# Human Amylin in the Presence of SARS-COV-2 Protein Fragments

**DOI:** 10.1101/2023.01.30.526275

**Authors:** Andrew D. Chesney, Buddhadev Maiti, Ulrich H. E. Hansmann

## Abstract

Covid-19 can lead to the onset of type-II diabetes which is associated with aggregation of islet amyloid polypeptides, also called amylin. Using molecular dynamics simulations, we investigate how the equilibrium, between amylin monomers in its functional form and fibrils associated with diabetes, is altered in presence of SARS-COV-2 protein fragments. For this purpose, we study the interaction between the fragment SFYVYSRVK of the Envelope protein or the fragment FKNIDGYFKI of the Spike protein with the monomer and two amylin fibril models. Our results are compared with earlier work studying such interactions for two different proteins.

## Introduction

SARS-COV-2 infections affect not only the respiratory system but a multitude of organs in the human body.^[1–4]^ However, the underlying mechanism for the resulting broad spectrum of symptoms and complications of COVID-19 are not always understood. For instance, diabetes patients have often more severe symptoms of COVID-19, but SARS-COV-2 infections may also trigger onset of diabetes.^[5–8]^ One possible cause for disease symptoms in individuals with type-II diabetes is aggregation of islet amyloid polypeptides (IAPP, also known as amylin). Amylin aggregation (or more general the onset of type-II diabetes) can be caused by various factors, for instance, inflammation induced by infections. However, one can speculate that amylin aggregation is also triggered directly by SARS-COV-2 protein fragments. In previous work, we have presented evidence for such a mechanism for Serum Amyloid A^[9]^ and α-synuclein^[10]^. However, amylin may be affected differently by interactions with viral proteins since it is more stable than these two proteins, being about 65% helical and stabilized by a disulfide bridge. For this reason, we investigate in this study whether interactions with SARS-COV-2 protein fragments also alter amylin amyloid formation, and how this effect differs from the one seen in our previous studies. For this purpose, we use all-atom molecular dynamics simulations to investigate the effect of two virus protein fragments (one, SK9, from the Envelope protein, and another, FI10, from the Spike protein, see **Figure 1a** and **Figure 1b**) on amylin monomers (**Figure 1c**) and two fibril models (**Figure 1d** and **Figure 1e**). Unlike in this earlier work on Serum Amyloid A^[9]^ and α-synuclein^[10]^ we do not see a shift in the ensemble of monomers toward more aggregation prone conformations, but rather protection of the native conformation. However, we observe stabilization of fibril structures that not only depends on the viral protein fragments but also on the amylin fibril geometry.

**Figure 1:**
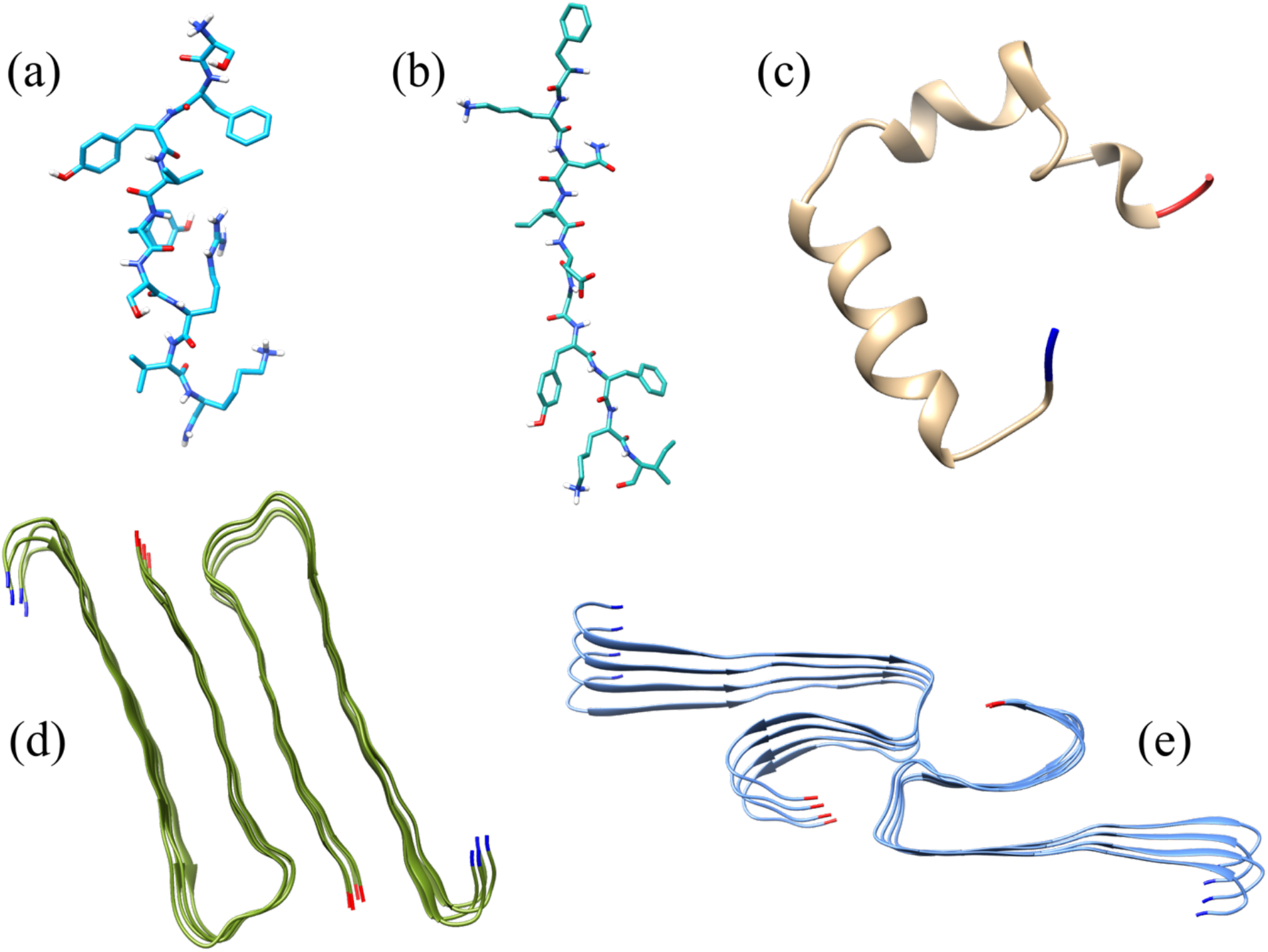
Licorice representation of the two SARS-COV-2 protein fragments used in this study: (a) SK9; (b) FI10. Ribbon representations of the (c) amylin monomer and (d, e) two fibril models are also shown. The N- and C-terminals are colored in blue and red, respectively.

## Results

### 1) MONOMER SIMULATIONS

We start our investigation into the effect of SARS-COV-2 protein fragments on amylin aggregation by first looking into the changes of the ensemble of monomer conformations induced by viral proteins. In our previous work we have demonstrated that presence of the nine-residue-long C-terminal fragment SK9 (SFYVYSRVK) of the SARS-COV-2 Envelope protein shifts the ensemble of SAA^[9]^ and αS^[10]^ monomers toward more aggregation-prone conformations, increasing in this way the probability for amyloid formation. The starting point for our present simulations is the amylin monomer model with PDB-ID: 2L86^[11]^ in complex with SK9. By comparing two different programs (AutoDock Vina and HADDOCK) to identify potential binding sites^[12–14]^ we come up with two distinct sets of trajectories. This allows us to probe the effect of the initial binding site on our results. A third set of trajectories was started from a configuration where the Spike protein fragment FI10 (FKNIDGYFKI) binds to the amylin monomer. This fragment, cleaved by the enzyme neutrophil elastase, which is released from neutrophils during acute inflammation, is unique for SARS-COV-2 and has been shown to form amyloids *in vitro*.^[15]^ Simulations of SK9 or FI10 interacting with amylin are compared with control simulations where the viral protein fragments are absent. PDB-files of start and final configurations of all runs are available as **supplemental material**.

In **Figure 2a**, we show the number of original binding contacts (contacts that exist at start) as function of time. Initially, SK9 has in one set (binding site derived with Autodock Vina) 37 contacts with the amylin monomer, 35 in the other set (binding site found with HADDOCK); and FI10 forms 38 contacts with the amylin monomer. In order to compare the various trajectories, we normalize the number of contacts for each system to a value of one at start. The log-log-plot in the inset shows that the number of contacts decreases by a power-law, with the exponent 0.34(1) for SK9 and of about 0.5 for FI10. These exponents indicate a diffusive motion of the viral protein fragments in relation to the amylin monomer, that in the case of SK9 is slower than in normal diffusion, and normal (i.e., random-walk-like) for FI10. Note, however, that in the process of this diffusive motion many of the disappearing original contacts are replaced by contacts not seen in the start conformation. As a consequence, the total number of contacts between the viral protein fragment and amylin decreases much slower, see **Figure 2b**, staying for SK9 close to 30%-40% of the initial number. For FI10 the number of contacts stays even at 60% -70%. The center-of-mass distance between both molecules is around 5 Å -20 Å, but occasionally increases up to 50 Å, see **Supplemental Figure SF2**. Hence, most of the time, the viral protein fragment stays attached to the amylin monomer, but occasionally it separates transiently. Together with our binding free energy estimates, the above observations indicate stable binding between the viral protein fragment and amylin. For SK9 we find a value of -21(3) kJ/mol for the trajectories starting from the Autodock Vina generated binding site, and a value of -22(5) kJ/mol for trajectories with HADDOCK generated initial binding site. These values are calculated taking the last 800 ns of the trajectories into account, and only decrease to -18.2 (3) kJ/mol and -18(5) kJ/mol, when only the last 200 ns are considered. Hence, the binding of SK9 to amylin depends little on the binding site and decreases only slowly. For FI10 we find values of -22(2) kJ/mol taking the last 800 ns into account, and -28(4) kJ/mol (when restricting the calculation to the last 200 ns), indicating that FI10 is binding more tightly with the amylin monomer than SK9. We remark that we observe the independence from the binding site, and the difference between SK9 and FI10, also when the binding energies are estimated directly from the start conformation, using Prodigy^[16]^. However, the obtained absolute values of the free energies, -45.6 (0) kJ/mol for SK9 docked with AutoDock Vina and -49.4 (7) kJ/mol for those docked with HADDOCK, and -41.6 (7) kJ/mol for FI10, are larger for Prodigy, which uses an approximation.

**Figure 2:**
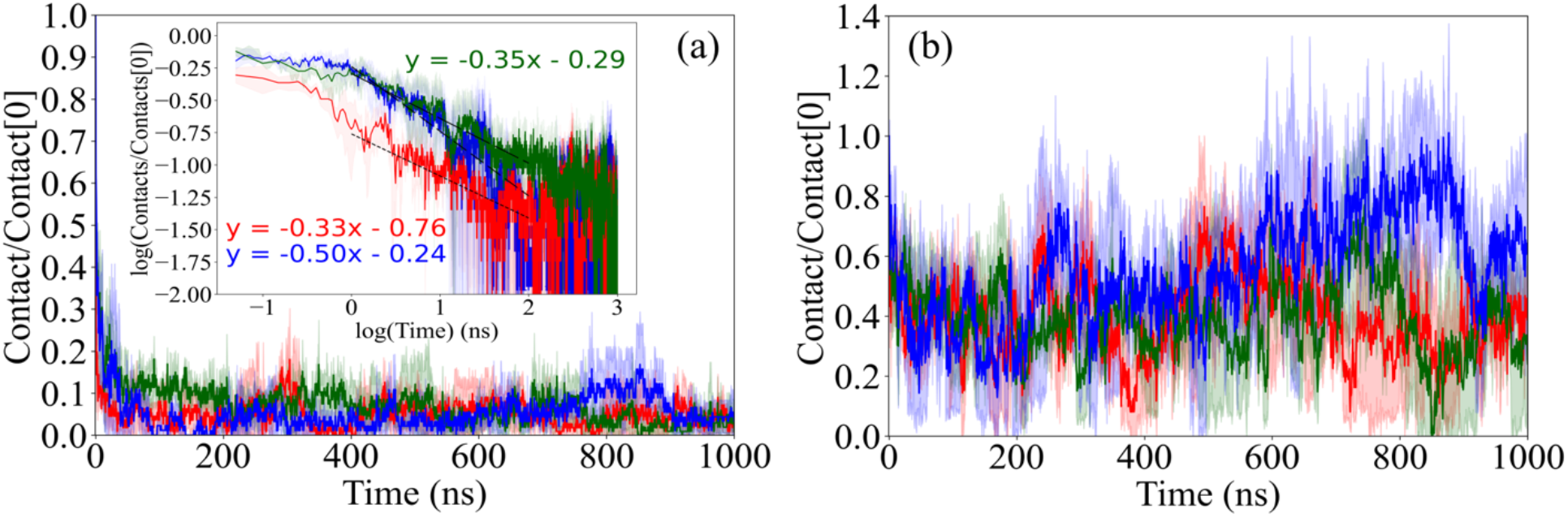
Averaged number of *original* contacts between SK9 and amylin as a function of time (a). The corresponding average number of *all* contacts (including newly formed ones) is shown in (b). Contacts are normalized to one for the respective start configurations. Red marks data from runs where the initial binding site of SK9 was found with AutoDock Vina, greens such where the binding site was found with HADDOCK. Blue marks data from runs where FI10 interacts with amylin.

Note that except for the terminal residues we find little difference in the frequencies that SK9 residues bind to amylin, with slightly higher values for the central Y5 and lower values for residues S6-S8 than for residues F2-V4. On average the binding probability of a SK9 residue is about 60(10)% for the last 800 ns, and 50(20)% when calculated only over the last 200 ns. Here, the terminal residues are excluded in the calculation of the binding probabilities. Similarly, we find for FI10 a binding probability of about 60(10)%, when measured over the last 800 ns, and 70(20)% when measured only over the last 200 ns. However, residues D5-I10 of FI10 have higher binding propensities than the first three residues (FKN), see **Supplemental Table ST1**. The frequencies are again independent from the initial binding site. Similarly, no clear pattern is seen in the amylin residues binding with SK9 or FI10, see **Supplemental Figure SF3**, reflecting the diffusive movement of the viral protein fragment in relation to the amylin monomer.

The effect of the binding of the viral protein fragment on the ensemble of amylin monomer conformations can be seen by comparing the simulations of amylin interacting with either SK9 or FI10 with the control simulations (where the viral protein fragments are absent). One example is the root-mean-square-deviation (RMSD) to the start conformation which we show as function of time in **Figure 3**. Shown are averages over all trajectories for each system. We have merged all trajectories of SK9 interacting with the amylin monomer as the data for SK9 differ little between the runs starting from binding sites generated with Autodock Vina and the ones generated by HADDOCK. After a rapid initial increase the RMSD stays in the control simulations, and for amylin interacting with either SK9 or FI10, almost constant over the whole length of the trajectory. However, while the RMSD values differ little between control and amylin in presence of either SK9 or FI10, visual inspection of the final configurations (**Figure 4**) shows differences in size and helicity to the control. For instance, the average radius of gyration of the amylin monomers is 11.4(2) Å for the SK9 and 12.9(1) Å for the FI10 simulations, which is higher than in the control 11.2(1) Å. A similar pattern is seen for the end-to-end distance, where we find 19(5) Å in the SK9 simulations, 26(5) Å in presence of FI10, and 20(1) Å for the control, see **Table 1**. While the average solvent accessible surface area (SASA) of the amylin monomer is similar in all cases, we find that the ratio of exposed surface of hydrophilic residues to that of hydrophobic residues is 0.86(2) in the simulations where FI10 interacts with amylin, which is comparable to the 0.85(3) measured in simulations where SK9 interacts with amylin, but both values are smaller than the value (0.875(5)) measured in the control simulations where neither of the viral protein fragments is present.

**Table 1:** Averages of various quantities measure in simulations of amylin monomer alone, or in presence of SK9 or FI10. Averages are over the last 200 or 800 ns, and over all respective trajectories. Contacts and binding free energies are between SK9 or FI10 and amylin. With the exception of the binding free energies are all quantities normalized to One at start.

| System | Time Interval | Total Contacts | Original Contacts | SASA | $R_g$ | Helicity | End-to-End Distance | $\Delta G_{\text{bind}}$ (kJ/mol) |
| --- | --- | --- | --- | --- | --- | --- | --- | --- |
| Control | 800ns | — | — | 0.96 (4) | 1.04 (5) | 0.6 (1) | 1.325 (5) | — |
|  | 200ns | — | — | 0.98 (7) | 1.1 (1) | 0.6 (2) | 1.27 (8) | — |
| w/ SK9 | 800ns | 0.39 (8) | 0.05 (2) | 0.96 (3) | 1.14 (6) | 0.79 (9) | 1.4 (3) | -21 (4) |
|  | 200ns | 0.33 (8) | 0.04 (1) | 0.96 (3) | 1.07 (6) | 0.68 (3) | 1.3 (4) | -18 (4) |
| w/ FI10 | 800ns | 0.60 (2) | 0.054 (7) | 1.00 (5) | 1.2 (1) | 0.79 (5) | 1.61 (9) | -22 (2) |
|  | 200ns | 0.70 (5) | 0.08 (3) | 1.03 (6) | 1.2 (1) | 0.72 (3) | 1.7 (3) | -28 (8) |

**Figure 3:**
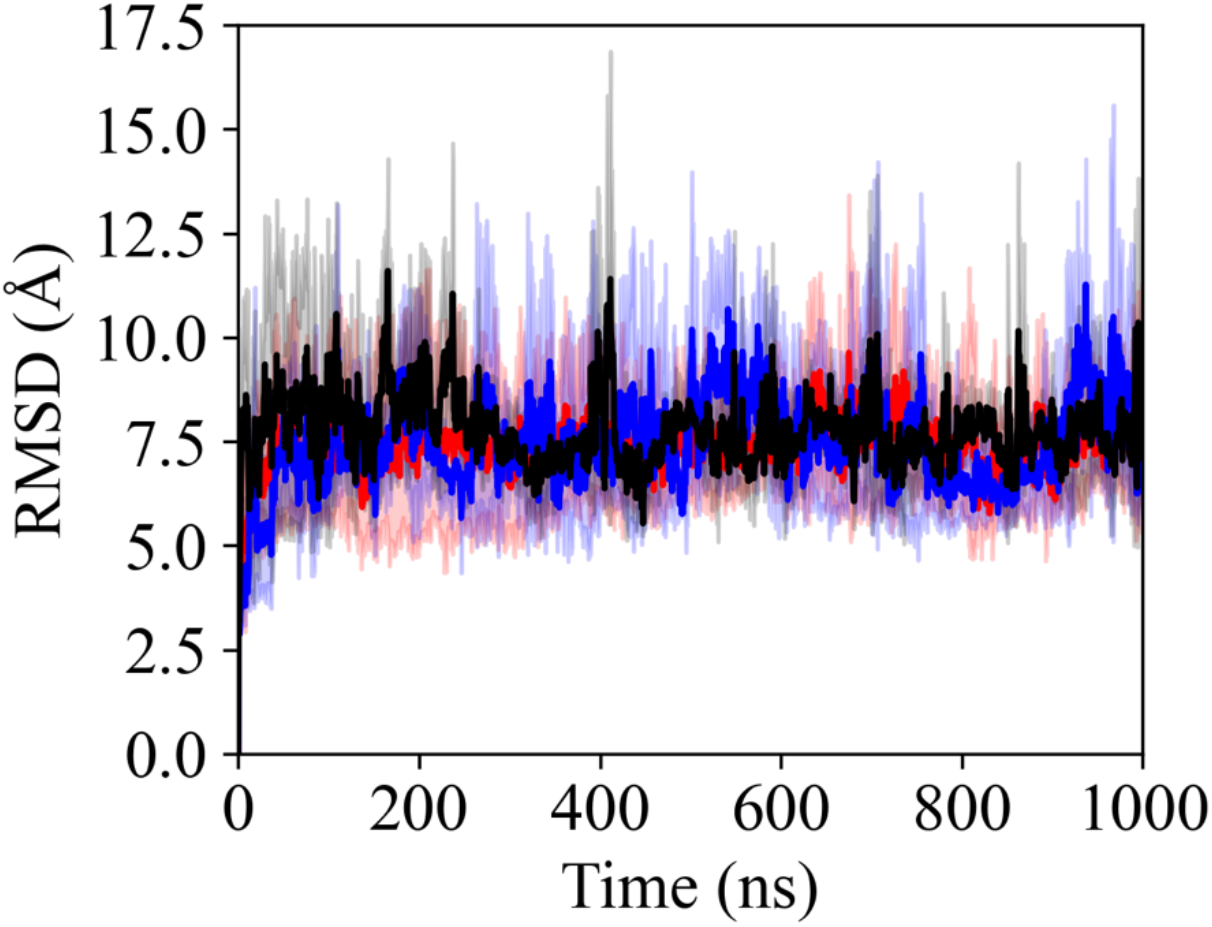
Average root-mean-square-deviation (RMSD) to the start conformation as function of time for amylin in presence of SK9 (red) or FI10 (blue), and in absence of SK9 and FI10 (black). The RMSD is evaluated over backbone atoms only. Averages are calculated over all trajectories for each system.

**Figure 4:**
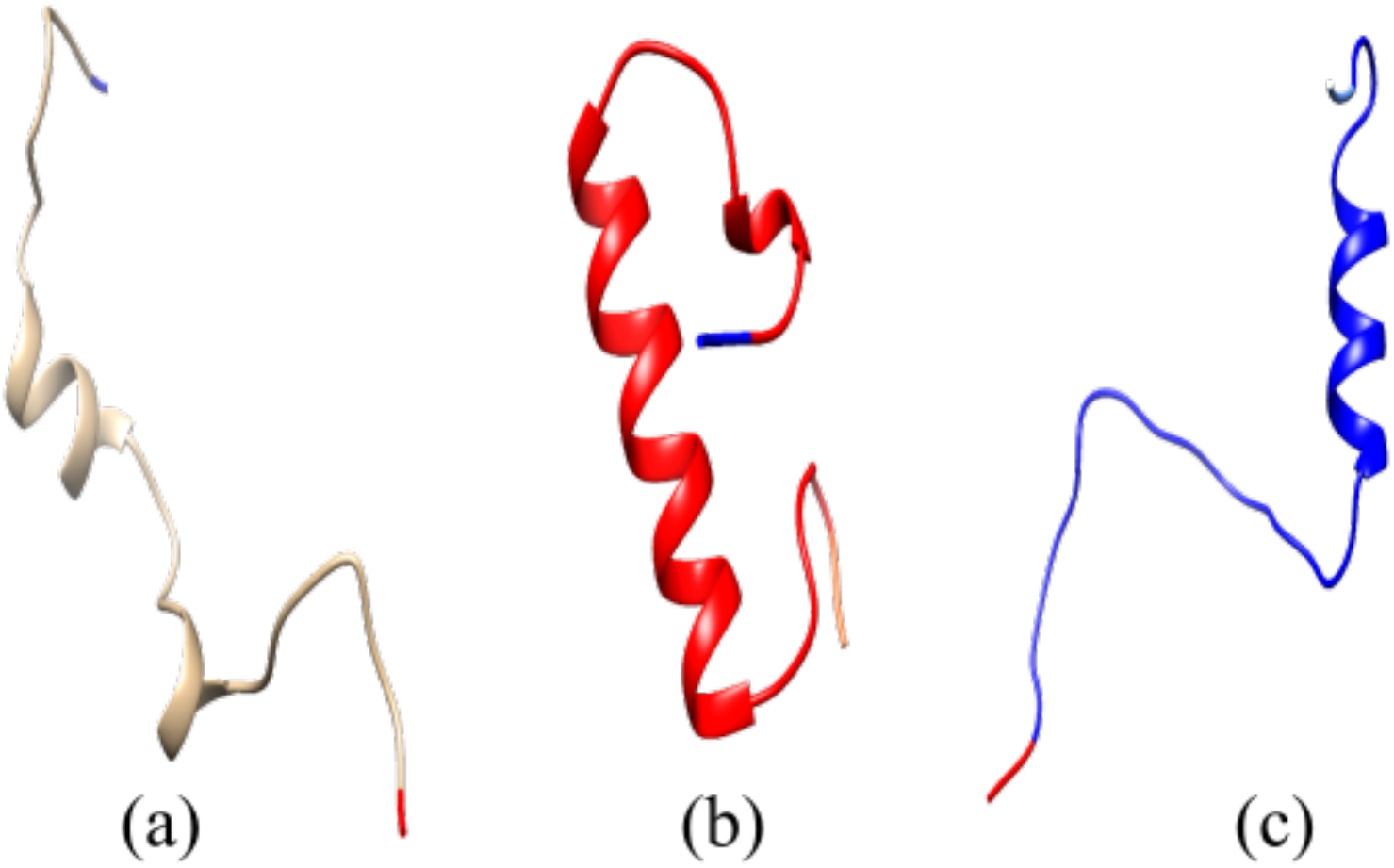
Representative snapshots of the final amylin conformations in absence (a), and presence of SK9 (b) or FI10 (c). The N- and C-terminals are colored in blue and red, respectively

The reason for the differences in these ratios, calculated from averages taken over the last 200 ns, can be seen in **Figure 5** where we show the root-mean-square-fluctuations (RMSF) of residues, evaluated over the last 200 and 800 ns. This quantity allows us to quantify the flexibility of residues. Hence, the lower values seen in presence of SK9 or FI10 indicate a stabilization of the monomer. Especially, we see in the control simulations that residues N22 - N31 have a higher flexibility than others, but this distinction is not seen in the simulations where the viral protein fragments are present. This is interesting as human amylin differs in the segment N22 - N31 from the less aggregation-prone rat amylin: rat amylin contains three proline residues between residues 20-29 that disrupt secondary structure due to structural hindrance and decreased flexibility of the protein backbone. As a result of these residues, also found in the amylin fibril inhibiting Pramlintide, rat amylin is less aggregation-prone and more soluble than human amylin. Hence, interactions with SK9 or FI10 mimic the protective effect of the rat mutations or in the amyloid inhibiting drug Pramlintide.

**Figure 5:**
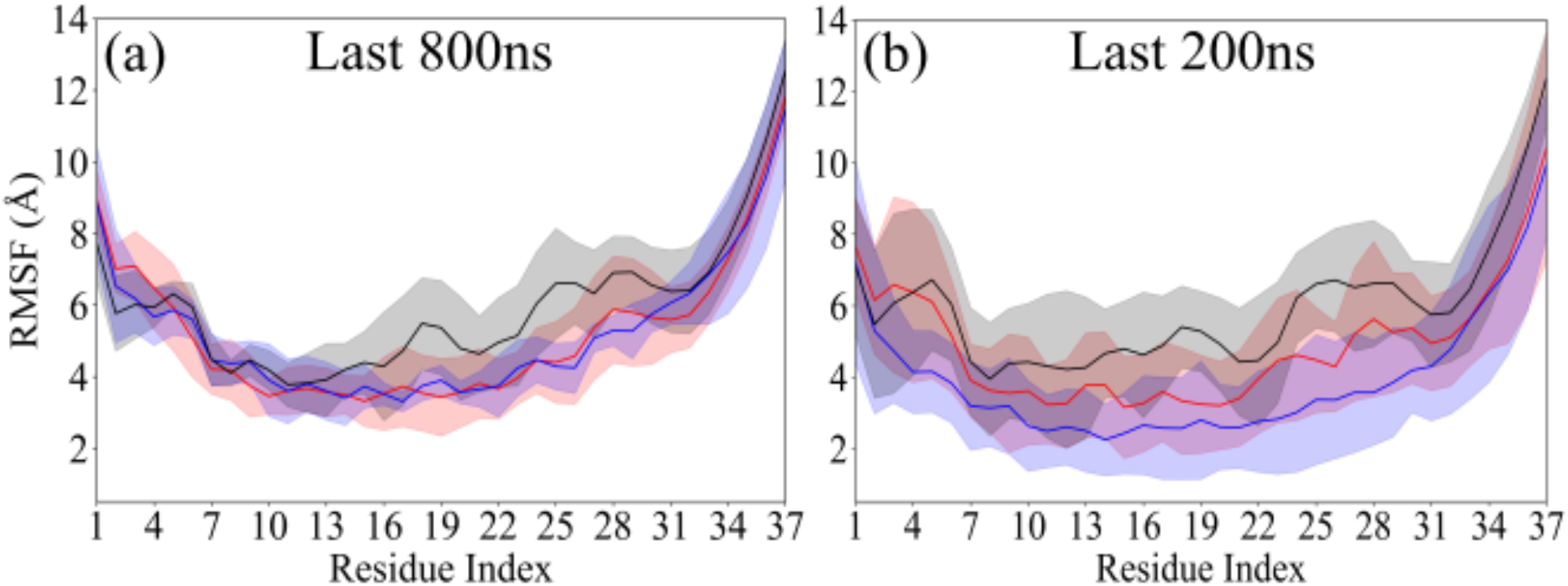
Residue-wise root-mean-square fluctuation of the amylin monomer in absence (black) or presence of SK9 (red) or FI10 (blue). Shaded regions mark the standard deviations of the shown quantities.

Indeed, we find that the average helicity, also shown in **Table 2**, stays marginally higher when amylin interacts with either SK9 (68(3)%) or FI10 (72(3)%) than in the control (60(20)%). However, these averages may be misleading. We show in **Figure 6** the residue-wise helicity measured over the last 200 ns in simulations with either SK9 or FI10 present, but subtracting the corresponding values measured in the control. In the presence of the viral protein fragments, residues F15 to S28 consistently have higher average helicity than observed in the control while residues A5 to N14 have a lower average helicity. This decrease of helicity for segment A5 to N14 is more pronounced for FI10, which at the same time also stabilizes helicity in residues L16 to T30 more than SK9. Residues S20 to S29 have been shown to form the primary amyloidogenic domain in amylin^[17]^ while the N-terminal residues (1-8) are not directly involved in fibril formation^[18]^. Hence, we conclude that both SK9 and FI10 are stabilizing the native monomer conformation of amylin, decreasing the chance of unfolding and seeding amyloids. This is unlike what we observed in our previous work where we found that SK9 shifted the ensemble of SAA^[9]^ and υS^[10]^ monomers toward more aggregation prone conformations, increasing in this way the probability for amyloid formation.

**Table 2:** Various quantities averaged over the last 100 ns of three trajectories in simulations of the amylin 2F4L fibril model in absence or presence of SK9.

|  | Control | w/ SK9 |
| --- | --- | --- |
| Total SASA (Å<sup>2</sup>) | 15554 (337) | 15251 (356) |
| Hydrophobic SASA (Å<sup>2</sup>) | 7782 (243) | 7623 (229) |
| Hydrophilic SASA (Å<sup>2</sup>) | 7773 (141) | 7628 (134) |
| Packing distance (Å) | 8.2 (8) | 8.1 (3) |
| Stacking distance (Å) | 2.9 (2) | 2.9 (2) |
| Packing contacts | 90 (11) | 92 (16) |
| Stacking contacts | 494 (26) | 492 (20) |
| Intrachain contacts | 1226 (6) | 1242 (3) |
| Packing hydrogen bonds | 6 (3) | 7 (3) |
| Stacking hydrogen bonds | 146 (5) | 142 (7) |
| Intrachain hydrogen bonds | 17 (1) | 16 (1) |
| Packing free energy (kJ/mol) | -189.2 (48.3) | -184.8 (77.0) |
| Stacking free energy (kJ/mol) | -251.1 (163.1) | -305.7 (113.7) |

**Figure 6:**
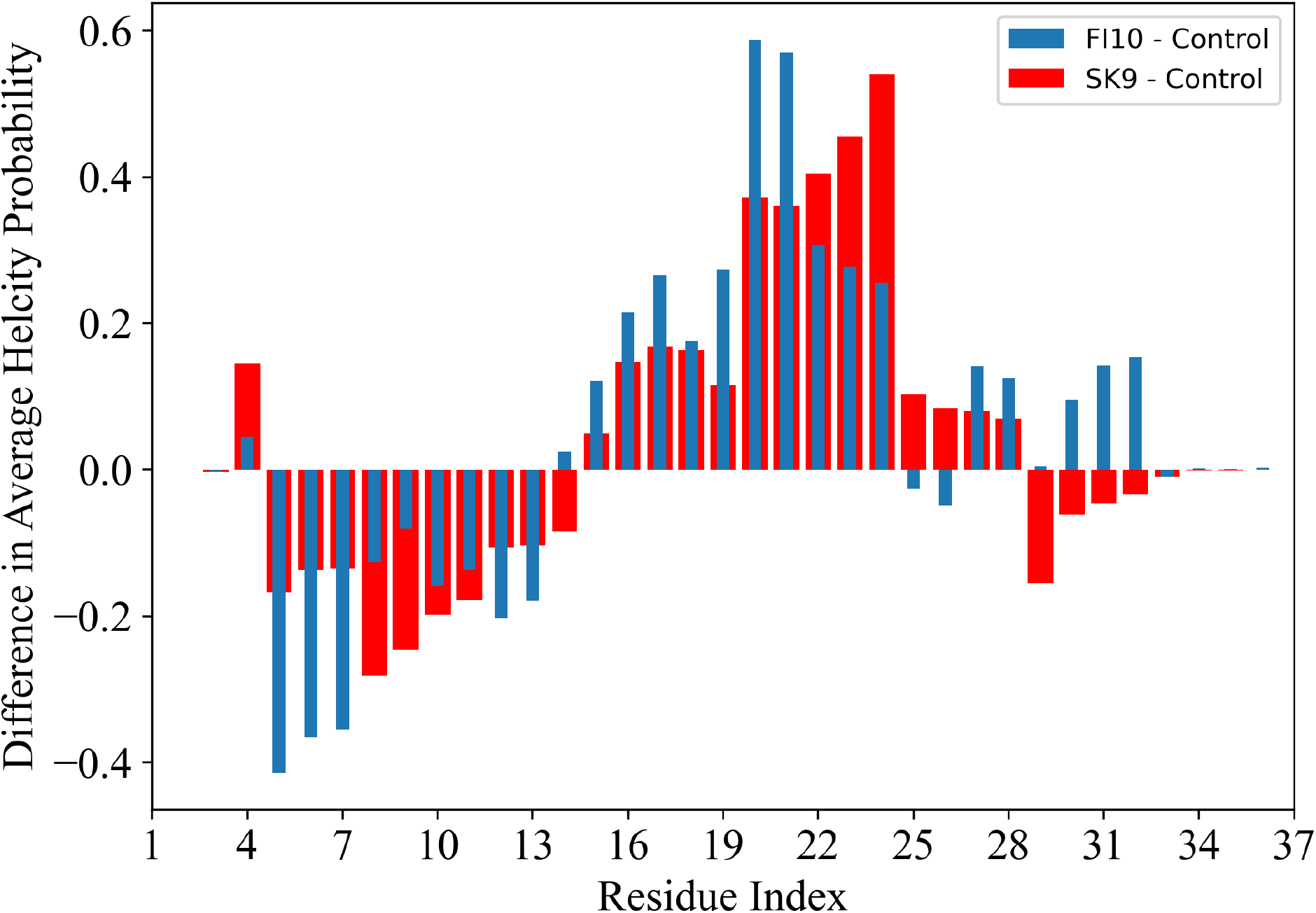
Residue-wise helicity of amylin monomer in presence of SK9 (red) and FI10 (blue) calculated over the last 200ns. Shown is the difference to the corresponding values in the control simulations where the viral protein fragments are absent.

### 2) FIBRIL SIMULATIONS

According to the Le Chatelier’s principle, the equilibrium between amylin in its functional form and aggregated in amyloids can be shifted not only by increasing the propensity of aggregation-prone monomer conformations, raising in this way the probability for amyloid formation, but also by stabilizing existing fibrils. Amyloids with more stable fibrillar structures have been shown to have higher rates of aggregation.^[19]^ In previous work^[9,10]^ we observed such stabilizing effect for the SK9 fragment on SAA and αS fibrils. Hence, we have also probed the change in stability of amylin fibrils introduced by interactions with virus protein fragments. Unfortunately, only few models of amylin fibrils have been resolved and published in the PDB. Because of our familiarity with the fibril model put forward by Eisenberg and co-workers^[20]^, which we have studied in a different context in previous work ^[21]^, we chose again this model, called by us 2F4L. PDB-files of start and final configurations are available as **supplemental material**.

At the start of the three trajectories, the viral SK9 peptides form 257 contacts with the 2F4L fibril, about 32 contacts per chain. This number decreases rapidly over the first nanosecond, and more slowly afterwards, but stays constant after 100 ns at 109(6) contacts, about 42% of the original number of contacts, i.e., on average 14(1) contacts per chain, leading to a binding free energy of -17(9) kJ/mol between an amylin and a SK9 peptide. Note that not all the observed contacts appear in the start configuration: the SK9 fragment is not tethered to the fibril and new contacts are formed as old original ones dissolve.

We show in **Figure 7a** the frequency with of amylin residues having a contact with SK9 as measured over the last 100 ns. The main binding regions include residues N14-S19 (centered around F15) in the outer layers of the protofibrils, and residues F23 and Y37 on the edge of the protofibril layers. Residues N14-S19 make hydrophobic contacts and π-π stacking interactions with SK9 to hold the SK9 close to protofibril layers, stabilizing their stacking. On the other hand, the residues F23 and Y37 form hydrogen bonds and hydrophobic contacts with SK9 that stabilize the packing of the two protofibrils. Hence, the effect of SK9 binding to the amylin chains is a slight stabilization of the fibril. This can be seen in **Figure 8a** where we compare the residue-wise root-mean-square fluctuation of the amylin fibril 2F4L bound to SK9 with that of the control (2F4L without SK9 present). This quantity is a measure for the flexibility of a residue, and while the overall form of the RMSF curves are similar, residues in the 2F4L fibril bound to SK9 have consistently lower RMSF values, i.e., are less flexible.

**Figure 7:**
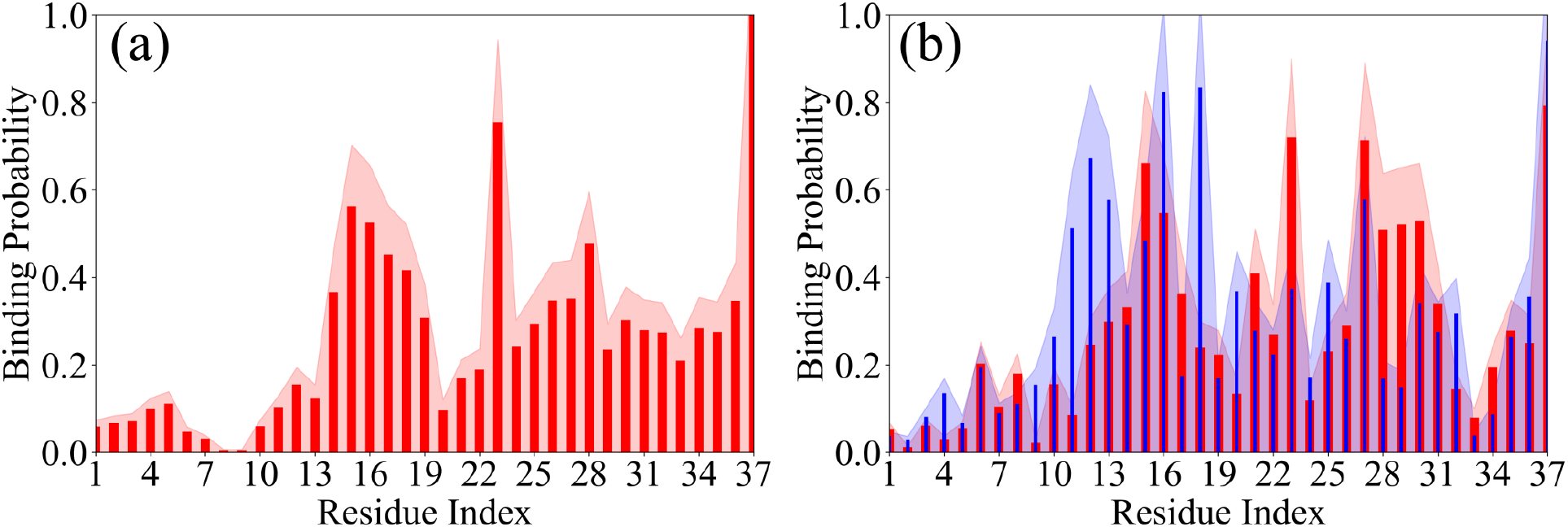
Residue-wise frequency of contacts of (a) the fibril model 2F4L with SK9, and (b) the fibril model 6ZRF, with either SK9 (red) or FI10 (blue). Shaded regions mark the standard deviation of the averages.

**Figure 8:**
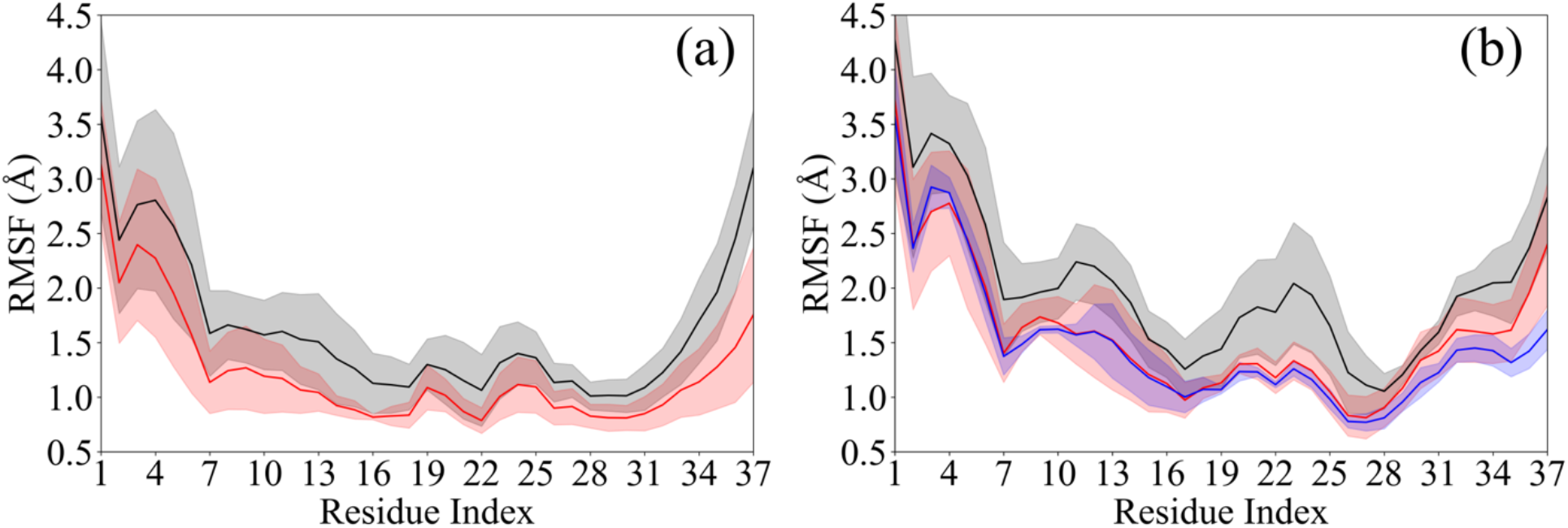
Residue-wise RMSF for (a) the fibril model 2F4L and (b) the fibril model 6ZRF. The black curve marks data from the simulation in absence of viral protein fragments, while the red curve show the results obtained when SK9 is present, and the blue curve data for when FI10 is interacting with the amylin fibrils. Shaded regions mark the standard deviation of the averages.

Consequently, the final configurations are less disturbed than in the control, see the insets of **Figure 9a** which shows as a function of time the root-mean-square deviation (RMSD) to the start conformation, taken over all backbone atoms of all chains. Note that we consider for the calculation of the RMSD only residues 8-37 as the first seven residues are disordered. While the differences to the control are small, they show larger stability of 2F4L bound to SK9 than for the fibril in absence of SK9. This can also be seen by visual inspection of the final conformations shown in the insets. Consistent with the RMSF plots, this stabilization is mainly due to the reduction of flexibility for the first N-terminal residues which in the control appear to disrupt the stacking of the N-terminal β-sheets.

**Figure 9:**
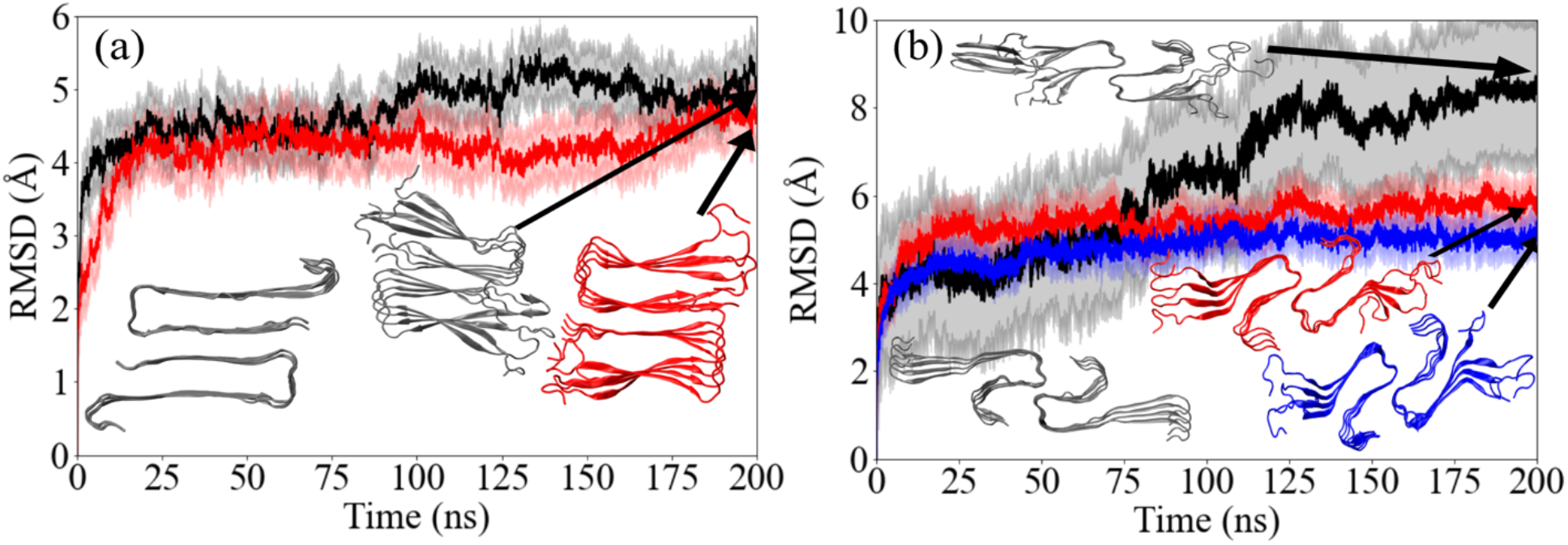
RMSD to the start configuration as function of time for (a) the fibril model 2F4L and the fibril model 6ZRF. The RMSD is calculated over all backbone atoms in residues 8-37 of all chains in the respective fibril model. Black curves are from the control simulations while red curves are from data measured in simulations where SK9 is present, and similarly, blue curves for the case of FI10 interacting with the amylin fibril. Shaded regions mark the standard deviation of the averages. The start configuration and representative final configurations are shown as insets.

However, these visual impressions are difficult to quantify. For instance, the solvent accessible surface area (SASA), in **Table 2**, is 15251(356) Å^2^ when measured in the simulations with SK9 which is smaller than when measured in the absence of SK9, 15554(337) Å^2^. The difference between the two averages is small and within the standard deviations, but the difference becomes more pronounced when excluding the disordered first seven residues from the SASA calculation, leading to 11445 (254) Å^2^ in presence of SK9 versus 11853(227) Å^2^ in the control. The reduction in exposed surface area in the complex is slightly higher for hydrophobic residues than for hydrophilic residues, but no clear picture emerges. Little difference is also seen for the average distance between the two folds (8.1(3) Å in presence of SK9 vs 8.2(8) Å for the control), and the averages distance between layers is with 2.9(2) Å about the same in presence and absence of SK9. Similarly, neither our MMPBSA approximations of the packing and stacking free energies nor the measured numbers of stacking or packing contacts or hydrogen bonds in **Table 2** give a clear signal explaining the observed weak stabilization.

Correspondingly, we do not see a difference in the frequency of the S29-S29 hydrogen bonds found in earlier work by us to be crucial for the packing of the fibril, or for the N31-N31 bond important for stacking^[21]^. However, we find the frequency of the F23-Y37 packing hydrogen bonds raised from 18% to 38%, and the distribution of packing hydrogen bond is tighter in presence of SK9. While in the simulations of the control more than 20 types of hydrogen bonds contribute at least 1% to the measured packing hydrogen bonds, only seven types of hydrogen bonds do this in the simulations where SK9 is present. Hence, while the number of packing hydrogen bonds changes little in presence of SK9, the bonds are less transient when SK9 interacts with the fibril. The more focused packing can be also seen in the map of packing contacts in **Supplemental Figure SF4a-c**.

The effect is less obvious for the stacking or intrachain hydrogen bonds. However, we observe an increase in contacts involving the N-terminal residues T4-L16, while such involving the C-terminal residues S20-Y37 decrease or shift more to off-diagonal contacts, see **Supplemental Figure SF4d-f**. This is consistent with the observed stabilization of the N-termini of the chains. As we find a slight increase in the number of intrachain contacts when SK9 is interacting with the 2F4L amylin fibril, increasing from 153(1) to 155(1), we conjecture that the main reason for the weak stabilization of this fibril geometry by SK9 is due to stabilization of the fold-geometry of the chains.

A characteristic of amyloids is the polymorphism of the resulting fibrils, and the virus proteins may change the amylin fibril stability differentially depending on the polymorph. Hence, we also evaluated the effect of SK9 on a second fibril model, resolved in 2020 by Gallardo et. al^[22]^ and deposited in the PDB under identifier 6ZRF. Simulations of this fibril model were also chosen by us to compare the effect of SK9 on fibril stability with that of the FI10 virus protein fragment. PDB-files of start and final configurations are again available as **supplemental material**.

The SK9 segments form 238 contacts (about 30 contacts per chain) initially with amylin chains in the 6ZRF fibril model, which is slightly less than in the 2F4L fibril. Again, these contacts rapidly decrease in the first 10 ns before the number of contacts stabilizes to 97(16) (about 40% of the initial number of contacts) over the last 100 ns. The binding free energy between SK9 and the 6ZRF amylin model is -31.4(4.4) kJ/mol, which is larger in magnitude than the 2F4L fibril. However, contacts are formed with similar residues as in the case of the 2F4L fibril, see **Figure 7b**: residues N14-V17 (centered around F15), N21, F23, L27 to T30 (centered around L27), and with highest probability to Y37. As for the 2F4L fibril these contacts result mostly from hydrophobic interactions, but also from π-π stacking (involving residues F15, F23, L27, S29 and Y37), or transiently formed hydrogen bonds (N14@O-SK9:Y5@HN, L16@O-SK9:Y3@HN, F23@O-SK9:Y3@HN, L27@O-SK9:Y3@HN and Y37@O-SK9:S1@H3). The net-effect is again a stabilization of the amylin fibril, see the reduced flexibility of residues in the RMSF plot of **Figure 8b** and a reduction in RMSD compared to the control that is more pronounced than for the 2F4L fibril geometry, see **Figure 9b**. For residues N14-V17, L27-T30 and Y37, increase these contacts with SK9 the frequency of stacking contacts between successive layers when compared to the control, see, for instance, in **Supplementary Figure SF6 (f)** residue N14 forming hydrogen bonds between its side chain carboxamide and other N14 residues in the protofibril. On the other hand, the interaction of SK9 with residues N21 and F23 strengthen the packing interactions between the two protofibrils. Residue F23, interacts with the hydrophobic core formed by residues L27-T30 and Y37 on the opposite protofibril which in turn brings the protofibrils closer together resulting in tighter packing. That SK9 stabilizes mainly packing of the protofibrils, avoiding the sliding of the protofibrils sometimes seen in the control simulations, is also confirmed by visual inspection of the final configurations, some of them shown as insets in **Figure 9b**.

As a result of this stabilization the reduction in solvent accessible surface area (SASA), resulting from interaction with the SK9 peptides, is more pronounced for the 6ZRF amylin fibril than for the 2F4L fibril model. This is also observed even when including the first seven residues, see **Table 3**, but again with little difference between hydrophobic and hydrophilic residues. Unlike for the 2F4L fibril, we also find a pronounced reduction in the packing distance (4.8(0.4) Å versus 5.9(1.4) Å in the control), which, however, is not seen in the distance between layers (3.0(0.2) Å versus 2.9(0.2) Å in the control), see **Table 3**. The differences in SASA values are consistent with a change in packing free energy of ΔΔG = -44.9 (49.1) kJ/mol between the two protofibrils, and ΔΔG = -41.5 (34.4) kJ/mol in stacking free energy between two layers, see **Table 3**. The lower binding free energies result from an increase in the number of packing contacts and layer-wise stacking contacts and hydrogen bonds, see **Table 3**. But note also the additional intra chain hydrogen bond observed in presence of SK9 that further stabilizes the amylin chain folds. As for the 2F4L fibril model, we find that presence of SK9 leads to a decrease in the number of different residue pairs that form packing hydrogen bonds: 15 such pairs contribute at least 1% in the control, while in presence of SK9 the number is reduced to five. The F23-A25 bond is dominant in both cases and contributes to about 70% of the packing hydrogen bonds. Interestingly, instead of the many kinds of transiently formed packing hydrogen bonds in the control, we find in the presence of SK9 a strong presence of N21-Y37 pairs. These pairs contribute 19% of the packing hydrogen bonds (instead of 2% in the control), and almost 90% of packing hydrogen bonds are of this type. As a consequence, the map of packing contacts in **Supplemental Figure SF5b** is more focused on contacts between residues in segment S20-A25 on one fold with residues N35-Y37 on the opposite fold than seen for the control (see **Supplemental Figure SF5a**, with the difference shown in **SF5c)**. Similarly, the frequency of stacking contacts, especially such involving residues A8-N14 also increases with the binding of SK9, see **Supplemental Figure SF5d-e**. These interactions stabilized the N-terminal β-sheets however, the effect is less pronounced than for the packing and goes with a free energy raising twisting of these sheets. Hence, the increased frequency of packing and stacking contacts is consistent with the MMGBSA estimates which indicate that in presence of SK9 both elongation and packing are favored more than in the control, with the effect slightly more pronounced for the packing of the protofibrils.

**Table 3:** Various quantities averaged over the last 100 ns of three trajectories in simulations of the amylin 6ZRF fibril model in absence or presence of SK9 or FI10.

|  | Control | w/ SK9 | w/ FI10 |
| --- | --- | --- | --- |
| Total SASA ($\text{\AA}^2$) | 12779 (112) | 12040 (267) | 11861 (284) |
| Hydrophobic SASA ($\text{\AA}^2$) | 6451 (160) | 6090 (85) | 6063 (263) |
| Hydrophilic SASA ($\text{\AA}^2$) | 6328 (68) | 5950 (265) | 5798 (129) |
| Packing distance ($\text{\AA}$) | 6 (1) | 4.8 (4) | 4.4 (1) |
| Stacking distance ($\text{\AA}$) | 2.9 (2) | 3.0 (2) | 2.9 (2) |
| Packing contacts | 54 (4) | 59 (5) | 62 (2) |
| Stacking contacts | 503 (10) | 537 (14) | 525 (13) |
| Intrachain contacts | 1254 (2) | 1255 (22) | 1256 (13) |
| Packing hydrogen bonds | 3 (1) | 2 (1) | 2 (1) |
| Stacking hydrogen bonds | 134 (4) | 139 (5) | 143 (7) |
| Intrachain hydrogen bonds | 22 (0) | 24 (1) | 25 (2) |
| Packing free energy (kJ/mol) | -152.6 (57.2) | -197.5 (39.3) | -222.6 (24.2) |
| Stacking free energy (kJ/mol) | -620.1 (38.5) | -661.6 (29.6) | -624.3 (50.9) |

Comparing the effect of FI10 with SK9 on the 6ZRF fibril, we note that the FI10 segments form 229 contacts (29 contacts per amylin chain) in the start configuration, which is fewer initial contacts than SK9. However, these contacts are more stable, and on average 139(6) contacts (about 61% of the original number) are observed over the last 100 ns, leading to a binding free energy between FI10 and amylin of -29 (3) kJ/mol that is only marginal lower than for SK9. The residue-wise distribution of contacts in **Figure 7b** is broader than for SK9. Binding sites such as R11-A13 (centered around L12) and H18 are observed more frequently while F23 and L27 have less pronounced peaks. Residue Y37 has an even higher binding probability to FI10 than to SK9. These contacts are again formed by hydrophobic interactions, χ-χ stacking and transiently formed hydrogen bonds (with N14@O-SK9:Y5@HN, L16@O-SK9:Y3@HN and L27@O- SK9:Y3@HN; and L16@O-SK9:Y3@HN replaced by L16@O-FI10:K2@HN, F23@O- SK9:Y3@HN by F23@O-FI10:G6@HN, and Y37@O-SK9:S1@H3 by two new ones: Y37@O- FI10:F1@HN and Y37@HN-FI10:F1@O). Formation of the additional hydrogen bond leads to the larger binding probability for residue Y37. FI10, unlike SK9, has a negatively charged residue that may reduce the flexibility of N-terminal residues by interacting with the positively charged R11. Similarly, interaction of the negatively charged residue on FI10 with H18 could reduce fibril destabilizing stress on the chain geometry appearing when the histidine is in the charged state.

Overall, we see a stronger interaction between FI10 and the fibril than between SK9 and the fibril. The RMSF plot of **Figure 8b** indicates that the resulting reduction in residue flexibility, when compared to the control, is slightly larger than for SK9, indicating larger stabilization of the fibril. This observation is supported by **Figure 9b** where the RMSD is lower than for SK9 binding to the fibril. Visual inspections of the final configuration, shown as inset of the figure, suggests that similar to SK9, the effect of FI10 on fibril stability is again mainly by stabilizing packing, and no sliding of the protofibrils was observed.

Consistent with this assumption is that the reduction in solvent accessible surface area upon interaction with the viral protein fragment is about 180 Å^2^ larger for FI10 than for SK9, when ignoring the disordered first seven residues. The additional reduction of SASA results mostly from hydrophilic residues, see **Table 3**. Packing distance (4.4 (1) Å) and the inter-layer distances (2.9 (2) Å) are likewise smaller than in the case of SK9, indicating that the larger stabilization of the amylin fibril by FI10, when compared to SK9, results from increased packing and stacking interactions. This can be also seen from our MMPBSA approximations of packing and stacking free energies which shows that packing of the protofibrils in the presence of FI10 is more favorable by ΔΔG = -70.0 (43.9) kJ/mol, and the stacking of layers by ΔΔG = -3.6(45.2) kJ/mol per chain. The large standard deviation in the stacking free energy difference makes it difficult to compare these results with that of SK9, but it appears that, unlike for SK9, the stabilization of the fibril is mostly due to that of improving the packing of the protofibrils. This is consistent with the larger number of packing contacts, exceeding that seen in control or when SK9 interact with fibril. On the other hand, differences between SK9 and FI10 are marginal for the stacking contacts and hydrogen bonds, see **Table 3**. However, similar to SK9, presence of FI10 leads to a reduction in the number of packing hydrogen bond forming residue pairs from 15 in the control to four pairs, with the N21-Y37 pairs now contributing 23% of the packing hydrogen bonds, and the F23-A25 hydrogen bond contributing 72%. Hence, 95% of packing hydrogen bonds result from only two types of hydrogen bonds, see also the map of packing contacts in **Supplemental Figure SF6a-c**. While we also see, an increase in stacking contacts, involving residues A8-N14, see **Supplemental Figure SF6d-e**, the fibril-stabilizing effect is even more than for SK9 due to enhancing the packing of the two protofibrils.

How do our results depend on our simulation set up? For instance, both virus protein fragments have in our simulations a NH_3_^+^-group at the N-terminus and a CONH_2_-group at the C-terminus. This is to reduce electrostatic interactions between the opposite charges at the termini. In order to test how our results depend on the choice of the end group, we have also simulated a system where FI10 is not capped at the C-terminus, i.e., ends with a COO^-^ group. We compare in supplemental figures **SF7** the RMSD as function of time for the control and the amylin fibril interacting with the two differently capped FI10 peptides. In both cases, the RMSD is lower than for the control, and therefore the values do not seem to depend on the end-group. However, in the RMSF plot of **supplemental figure SF8** values of the FI10 peptide with uncapped C-terminus are higher than for the capped one, but still lower than the control. Other quantities such as the SASA, 12360(502) Å^2^, are closer to the values found for the control, 12779 (112) Å^2^, than for the fibril in presence of the capped FI10 peptides, 11861(284) Å^2^. Unlike for the capped peptide, the differences result mostly from hydrophilic residues. Only residue 8-37 were considered in calculating the above SASA values. When including the disordered first seven residues the values for control, 16848 (257) Å^2^, and the fibril in presence of the uncapped peptide, 16834(459) Å^2^, agree within the error bars, while still lower for the fibril in presence of the capped FI10, 16283(577) Å^2^. This may be because the uncapped peptide forms more compact configurations as its average end-to-end distance of 20.6(9) Å is smaller than the 23(1) Å of the capped FI10. Therefore, the uncapped peptide binds less tightly to the amylin fibril than the capped one. This can be seen with the number of contacts in the last 100 ns being only about 116 (6), i.e., 51% of the original number of contacts seen for the uncapped FI10 versus 139(6), i.e., 61% for the capped peptide. The residue-wise binding frequencies in **supplemental figure SF9** show a similar, but broader, distribution as for the capped peptide, with for the N-terminal residues K1-H18 less binding propensity. Correspondingly, we find that the difference in binding free energies, ΔΔG = -10.5 (2.3) kJ/mol, resulting from the different end groups, indicates less of an interaction between the uncapped virus protein fragments and the amylin fibril. Hence, we conclude that capping of the termini reduces self-interaction in the FI10, leading to more stretched conformations which in turn interact more strongly with the amylin fibril. This effect, however, does not seem to reduce the overall stabilization of the fibril.

## Conclusions

Using extensive molecular dynamics simulations, we have studied the effect of two virus protein fragments from the Envelope protein (SK9) and from the Spike protein (FI10) of SARS-COV-2 on the ensemble of amylin monomer conformations, and on the stability of amylin fibrils. Such interactions may be correlated with the observed onset of type-2 diabetes following COVID-19.^[5– 8]^ While both virus protein fragments bind to the amylin monomer structure, we do not find the induced shift toward more aggregation-prone conformations that we saw previously in our studies with Serum Amyloid A^[9]^ and α-synuclein^[10]^. Unlike these proteins amylin has, even as a monomer, a more defined structure, and the mostly helical fold appears to be stabilized by the presence of SK9 or FI10. Indications for such a protective effect seems to be seen also in experiments by the Huang Lab at HUKST that are currently analyzed (Jinqing Hiuang, private communication).

However, presence of amylin aggregates, proposed to cause symptoms of type-2 diabetes, can also be raised by stabilizing existing fibrils, shifting the equilibrium from functional monomers toward the toxic amyloids. We indeed see such stabilization of amylin fibrils by the viral protein fragments. Similar to what we found in our previous work, this stabilizing effect depends on the fibril model. Hence, as also observed by us earlier for α-synuclein^[10]^, interactions with the viral protein fragments may shift the equilibrium between various fibril polymorphs. Fibril stabilization is observed for both SK9 and FI10 and depends little on details of our simulation set up such as the capping of the viral protein fragments. The stabilization of fibril geometry is mainly by enhancing the packing of protofibrils and less by encouraging elongation of fibrils. Interestingly, the effect is stronger for FI10. This segment unique is for SARS-COV-2 and cleaved from the spike protein by the enzyme neutrophil elastase, released from neutrophils during acute inflammation. In vitro, the FI10 fragment forms amyloids.^[15]^ While not yet supported by experimental evidence, we speculate that SARS-COV-2 infections, often causing chronic inflammation, increase enzymatic cleavage of the spike protein, releasing in some cases amyloidogenic fragments such as FI10. These fragments could then cross-seed amylin fibrils which in turn may contribute to an onset of type-2 diabetes.

Comparing our results with earlier work we find as a common theme that small SARS-COV-2 protein fragments (as potentially derived by cleavage during infection-cause inflammation) can seed aggregation of amyloidogenic human proteins, but that this effect depends both on the viral fragment and the structure of the fibril polymorph.

## Materials and Methods

### System Preparation

We rely on molecular dynamics (MD) simulations to study the effect of SARS-COV-2 proteins on amylin aggregation. Our study focuses on the change in the conformational ensemble and the stability of amylin monomer and fibrils induced by presence of the nine-residue segment SFYVYSRVK (SK9) of the Envelope protein or the ten-residue segment FKNIDGYFKI (FI10) of the Spike protein from SARS-COV-2. The initial configuration of SK9 is the same as the one used in our previous work,^[9,10]^ and was derived from the model of the SARS-COV-2 Envelope protein deposited in the MOLSSI COVID-19 hub^[23]^. The N- and C-terminal of the SK9 segment are capped by a NH_3_^+^ and -CONH_2_ group, respectively. Following Nyström and Hammarström^[15]^ the initial configuration of the FI10 viral fragment was derived by extracting the residues F194- I203 from the structure of the SARS-COV-2 Spike protein (PDB ID: 6VXX), with the N- and C-terminal of the peptide capped again by a NH_3_^+^ and -CONH_2_ group. To compare the effect of terminal group of the small FI10 peptide on our results, we have also considered the case where the N- and C-termini were capped with a NH_3_^+^ and -COO^−^ group, respectively.

The initial conformation of the amylin monomer model has been taken from the Protein Data Bank (PDB). This structure (PDB ID: 2L86 ^[11]^) was resolved by solution NMR, and we add at the N- and C-terminals a NH_3_^+^ and a COO^−^ group respectively. Unfortunately, only a few models of amylin fibrils have been resolved and published in the PDB. Due to our familiarity with the fibril model put forward by Eisenberg and co-workers^[20]^, which we have studied in a different context in previous work ^[21]^, we again chose this model, but reduced the number of layers from five to four chains in each of the two proto-fibrils. This is because we expect that by decreasing the layers in this manner the stability of the fibril fragment will be lowered, and any potential effect resulting from presence of SK9 will be seen more clearly. Note that while we call this amylin fibril model 2F4L, this is not a PDB-ID but rather our shorthand notation for **T**wo-**F**olds-**F**our-**L**ayers. In order to be consistent with earlier work^[21]^, we add again at the N- and C- terminal a NH3^+^ and a COO^−^ group. The initial conformation of the other amylin fibril model was extracted from the Cryo-EM resolved model^[22]^ with PDB-ID: 6ZRF such that it has the same number of chains as the 2F4L fibril model. Note that we use for this second fibril model -COCH_3_ and -NHCH_3_ as end groups, allowing us to compare our control simulations (in absence of viral protein fragments) with the experiments described in Ref ^[24]^. The use of different end-groups is justified as our main interest is the comparison of simulations with and without SARS-COV-2 protein fragments.

Simulations starting from the amylin monomer and fibril models described above serve as control in our study. For investigating the effect of SARS-COV-2 protein fragments on amylin we have performed simulations that start from configurations where either SK9 or FI10 peptides bind to these models. These start configurations were generated using AutoDock Vina^[12]^ and HADDOCK^[13, 14]^ by docking the respective segments in a ratio of 1:1 with either monomer or fibrils. The resulting configurations are shown in **Supplemental Figure S1**. Note that the viral protein segments are not restricted to these initial positions on monomer or fibrils but can move freely throughout the simulations and even detach from the amylin.

### General Simulation Protocol

The set-up and simulation of all systems rely on the GROMACS 2018 and GROMACS 2022 package^[25]^. We use the CHARMM 36m all-atom force-field^[26]^ with TIP3P explicit water^[27]^ as implemented in the GROMACS package to describe the inter- and intramolecular interactions for the monomer and fibrils. This force-field and water model combination has been found in previous work performed in our group^[21,28]^ and in literature^[29–31]^ to be well-suited for simulations of fibrils and oligomers. Hydrogen atoms are added with the *pdb2gmx* module of the GROMACS suite^[25]^. The start configurations for all systems are put in the center of the cubic box, with at least 15 Å between the solute and the edge of the box. Periodic boundary conditions are employed. The systems are solvated with water molecules, and counterions are added to neutralize the system, with the Na^+^ and Cl^-^ ions at a physiological ion concentration of 150 mM NaCl. Both the number of water molecules and the total number of atoms are listed in **Table 4**. The energy of each system is minimized by steepest decent for up to 50,000 steps, and afterwards the system is equilibrated at 310K for 200 ps at constant volume and in an additional 200ps at constant pressure (1 atm), constraining the positions of heavy atoms with a force constant of 1000 kJ mol^-1^ nm^-2^.

**Table 4:** Simulated systems

| System Description | Atoms | Water Molecules | Independent Trajectories | Simulation Length (ns) | Total Sampling (ns) |
| --- | --- | --- | --- | --- | --- |
| **Monomer (PDB ID: 2L86)** |  |  |  |  |  |
| Control | 26,747 | 8,720 | 3 | 4,000 | 12,000 |
| w/ SK9 (AD-Vina) | 26,767 | 8,733 | 3 | 1,000 | 3,000 |
| w/ SK9 (HADDOCK) | 25,856 | 8,430 | 3 | 1,000 | 3,000 |
| w/ FI10 | 30,416 | 10,083 | 3 | 1,000 | 3,000 |
| **Fibril Model 2F4L** |  |  |  |  |  |
| Control | 118,169 | 37,884 | 3 | 200 | 600 |
| w/ SK9 | 120,148 | 38,090 | 3 | 200 | 600 |
| **Fibril Model 6ZRF** |  |  |  |  |  |
| Control | 149,710 | 48,356 | 3 | 200 | 600 |
| w/ SK9 | 268,411 | 87,361 | 3 | 200 | 600 |
| w/ FI10-CONH<sub>2</sub> | 268,423 | 87,373 | 3 | 200 | 600 |
| w/ FI10-COO<sup>-</sup> | 268,433 | 87,403 | 3 | 200 | 600 |

Starting from the above generated initial conformations (shown in Supplemental **Figure SF1**) simulations are performed at constant temperature (310 K) and pressure (1 atm). The temperature is controlled by a v-rescale thermostat^[32]^ with a coupling constant of 0.1 ps, and pressure is kept constant by using the Parrinello-Rahman barostat^[33]^ with a pressure relaxation time of 2 ps. By using the SETTLE algorithm^[34]^ to keep water molecules rigid, and restraining protein bonds involving hydrogen atoms to their equilibrium length with the LINCS algorithm^[35]^, we are able to use a time step of 2 fs for integrating the equations of motion. Because of the periodic boundary conditions, we compute the long-range electrostatic interactions with the particle-mesh Ewald (PME) technique, using a real-space cutoff of 12 Å and a Fourier grid spacing of 1.6 Å. Short-range van der Waal interactions are truncated at 12 Å, with smoothing starting at 10.5 Å. For each system, we follow three independent trajectories differing by their initial velocity distributions. The length of the various trajectories is also listed in **Table 4**.

### Trajectory Analysis

The molecular dynamics trajectories are analyzed with the GROMACS tools^[25]^, VMD^[36]^ and MDTraj software^[37]^. For visualization of trajectory conformations we use VMD^[36]^, and UCSF Chimera^[38]^. The root-mean-square deviation (RMSD), root-mean-square-fluctuation (RMSF), radius of gyration (RGY) and solvent accessible surface area (SASA) are calculated using GROMACS tools, for the latter quantity using a spherical probe of 1.4 Å radius. The residue-wise contact frequencies are calculated using VMD and MDTraj software, defining contacts by a cutoff 4.5 Å in the closest distance between heavy atoms in a residue pair. Hydrogen bonds are defined by a distance cutoff of 3.5 Å between the donor and acceptor atoms and an angle cutoff of 30^°^. The residue-wise secondary structure propensity is calculated in VMD by the STRIDE algorithm^[39]^.

In order to avoid the in our case prohibitive computational costs of free energy perturbation methods, thermodynamic integration, or similar approaches are binding free energies of the viral protein fragments to amylin fibrils approximated by the molecular mechanics Poisson-Boltzmann surface area (MM-PBSA) method as implemented in the GROMACS suite^[25,40]^. This approximation is not without problems, with one important drawback being that it neglects entropic contributions of the solute. As MM-PBSA is therefore not suitable for studying the binding of the viral protein fragments SK9 and FI10 to amylin monomers, we use for these systems instead the approach described in Bellaiche et. al^[41]^. This approach was used earlier to study inhibition of Aβ amyloid formation by small peptides^[42]^ and relies on calculating a dimensionless association constant from the distribution of bound and unbound conformations. For comparison, we also estimate binding free energies with the protein binding energy predictive model (PRODIGY) which is based on a simple linear regression of the number of interfacial contacts between residues in a protein-protein complex^[16]^.

## Supporting information

Folder with start and final configurations for all trajectories

## AUTHOR INFORMATION

### Notes

The authors declare no competing financial interest.

### Supporting Information

The Supporting Information is available free of charge at URL

## Acknowledgement

Our simulations were done using the SCHOONER cluster of the University of Oklahoma, XSEDE resources allocated under grant MCB160005 (National Science Foundation), and TACC resources allocated under grant under grant MCB20016 (National Science Foundation). We acknowledge financial support from the National Institutes of Health under grant GM120634. We thank Dr. Asis K. Jana for help on early stages of this project, mainly setting up and running the 2F4L fibril simulations and early parts of the trajectories of amylin monomers interacting with SK9.

## For Table of Contents Only

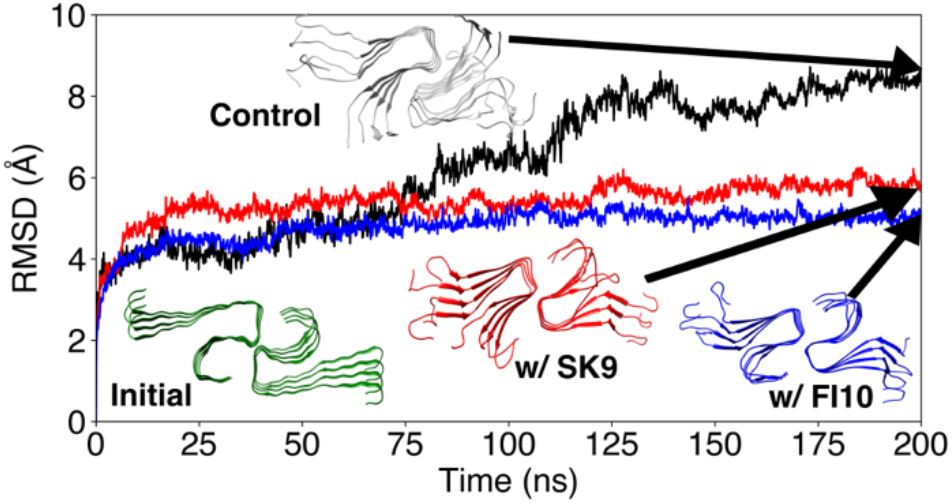

## Notes

### Competing Interest Statement

The authors have declared no competing interest.

## REFERENCES

(1) Shi, S.; Qin, M.; Shen, B.; Cai, Y.; Liu, T.; Yang, F.; Gong, W.; Liu, X.; Liang, J.; Zhao, Q.; Huang, H.; Yang, B.; Huang, C. Association of Cardiac Injury With Mortality in Hospitalized Patients With COVID-19 in Wuhan, China. JAMA Cardiol. 2020, 5 (7), 802. https://doi.org/10.1001/jamacardio.2020.0950.

(2) Sheraton, M.; Deo, N.; Kashyap, R.; Surani, S. A Review of Neurological Complications of COVID-19. Cureus 2020. https://doi.org/10.7759/cureus.8192.

(3) Su, H.; Yang, M.; Wan, C.; Yi, L.-X.; Tang, F.; Zhu, H.-Y.; Yi, F.; Yang, H.-C.; Fogo, A. B.; Nie, X.; Zhang, C. Renal Histopathological Analysis of 26 Postmortem Findings of Patients with COVID-19 in China. Kidney Int. 2020, 98 (1), 219–227. https://doi.org/10.1016/j.kint.2020.04.003.

(4) Morris, S. B.; Schwartz, N. G.; Patel, P.; Abbo, L.; Beauchamps, L.; Balan, S.; Lee, E. H.; Paneth-Pollak, R.; Geevarughese, A.; Lash, M. K.; Dorsinville, M. S.; Ballen, V.; Eiras, D. P.; Newton-Cheh, C.; Smith, E.; Robinson, S.; Stogsdill, P.; Lim, S.; Fox, S. E.; Richardson, G.; Hand, J.; Oliver, N. T.; Kofman, A.; Bryant, B.; Ende, Z.; Datta, D.; Belay, E.; Godfred-Cato, S. Case Series of Multisystem Inflammatory Syndrome in Adults Associated with SARS-CoV-2 Infection — United Kingdom and United States, March–August 2020. MMWR Morb. Mortal. Wkly. Rep. 2020, 69 (40), 1450–1456. https://doi.org/10.15585/mmwr.mm6940e1.

(5) Beliard, K. A.; Yau, M.; Wilkes, M.; Romero, C. J.; Wallach, E.; Rapaport, R. SARS-CoV-2 Infection Related Diabetes Mellitus. J. Endocr. Soc. 2021, 5 (Supplement_1), A397–A397. https://doi.org/10.1210/jendso/bvab048.808.

(6) Hollstein, T.; Schulte, D. M.; Schulz, J.; Glück, A.; Ziegler, A. G.; Bonifacio, E.; Wendorff, M.; Franke, A.; Schreiber, S.; Bornstein, S. R.; Laudes, M. Autoantibody-Negative Insulin-Dependent Diabetes Mellitus after SARS-CoV-2 Infection: A Case Report. Nat. Metab. 2020, 2 (10), 1021–1024. https://doi.org/10.1038/s42255-020-00281-8.

(7) Moreno, D. M.; Ramos, R. J. A.; Fernández, L. G.; Montenegro, A. M. R.; González, M. M.; Torrecilla, N. B.; Albarrán, O. G. Clinical/Biochemical Characteristics and Related Outcomes in People with New-onset Diabetes and COVID -19: Experience from a Single Centre. Pract. Diabetes 2022, 39 (6), 24–31. https://doi.org/10.1002/pdi.2426.

(8) Hayden, M. R. An Immediate and Long-Term Complication of COVID-19 May Be Type 2 Diabetes Mellitus: The Central Role of β-Cell Dysfunction, Apoptosis and Exploration of Possible Mechanisms. Cells 2020, 9 (11), 2475. https://doi.org/10.3390/cells9112475.

(9) Jana, A. K.; Greenwood, A. B.; Hansmann, U. H. E. Presence of a SARS-CoV-2 Protein Enhances Amyloid Formation of Serum Amyloid A. J. Phys. Chem. B 2021, 125 (32), 9155– 9167. https://doi.org/10.1021/acs.jpcb.1c04871.

(10) Jana, A. K.; Lander, C. W.; Chesney, A. D.; Hansmann, U. H. E. Effect of an Amyloidogenic SARS-COV-2 Protein Fragment on α-Synuclein Monomers and Fibrils. J. Phys. Chem. B 2022, 126 (20), 3648–3658. https://doi.org/10.1021/acs.jpcb.2c01254.

(11) Nanga, R. P. R.; Brender, J. R.; Vivekanandan, S.; Ramamoorthy, A. Structure and Membrane Orientation of IAPP in Its Natively Amidated Form at Physiological PH in a Membrane Environment. Biochim. Biophys. Acta BBA - Biomembr. 2011, 1808 (10), 2337–2342. https://doi.org/10.1016/j.bbamem.2011.06.012.

(12) Trott, O.; Olson, A. J. AutoDock Vina: Improving the Speed and Accuracy of Docking with a New Scoring Function, Efficient Optimization, and Multithreading. J. Comput. Chem. 2009, NA-NA. https://doi.org/10.1002/jcc.21334.

(13) van Zundert, G. C. P.; Rodrigues, J. P. G. L. M.; Trellet, M.; Schmitz, C.; Kastritis, P. L.; Karaca, E.; Melquiond, A. S. J.; van Dijk, M.; de Vries, S. J.; Bonvin, A. M. J. J. The HADDOCK2.2 Web Server: User-Friendly Integrative Modeling of Biomolecular Complexes. J. Mol. Biol. 2016, 428 (4), 720–725. https://doi.org/10.1016/j.jmb.2015.09.014.

(14) Honorato, R. V.; Koukos, P. I.; Jiménez-García, B.; Tsaregorodtsev, A.; Verlato, M.; Giachetti, A.; Rosato, A.; Bonvin, A. M. J. J. Structural Biology in the Clouds: The WeNMR-EOSC Ecosystem. Front. Mol. Biosci. 2021, 8, 729513. https://doi.org/10.3389/fmolb.2021.729513.

(15) Nyström, S.; Hammarström, P. Amyloidogenesis of SARS-CoV-2 Spike Protein; preprint; Biochemistry, 2021. https://doi.org/10.1101/2021.12.16.472920.

(16) Xue, L. C.; Rodrigues, J. P.; Kastritis, P. L.; Bonvin, A. M.; Vangone, A. PRODIGY: A Web Server for Predicting the Binding Affinity of Protein–Protein Complexes. Bioinformatics 2016, btw514. https://doi.org/10.1093/bioinformatics/btw514.

(17) Westermark, P.; Engström, U.; Johnson, K. H.; Westermark, G. T.; Betsholtz, C. Islet Amyloid Polypeptide: Pinpointing Amino Acid Residues Linked to Amyloid Fibril Formation. Proc. Natl. Acad. Sci. 1990, 87 (13), 5036–5040. https://doi.org/10.1073/pnas.87.13.5036.

(18) Jaikaran, E. T. A. S.; Higham, C. E.; Serpell, L. C.; Zurdo, J.; Gross, M.; Clark, A.; Fraser, P. E. Identification of a Novel Human Islet Amyloid Polypeptide β-Sheet Domain and Factors Influencing Fibrillogenesis. J. Mol. Biol. 2001, 308 (3), 515–525. https://doi.org/10.1006/jmbi.2001.4593.

(19) Thu, T. T. M.; Li, M. S. Protein Aggregation Rate Depends on Mechanical Stability of Fibrillar Structure. J. Chem. Phys. 2022, 157 (5), 055101. https://doi.org/10.1063/5.0088689.

(20) Wiltzius, J. J. W.; Sievers, S. A.; Sawaya, M. R.; Cascio, D.; Popov, D.; Riekel, C.; Eisenberg, D. Atomic Structure of the Cross-β Spine of Islet Amyloid Polypeptide (Amylin). Protein Sci. Publ. Protein Soc. 2008, 17 (9), 1467–1474. https://doi.org/10.1110/ps.036509.108.

(21) Pandey, P.; Nguyen, N.; Hansmann, U. H. E. D -Retro Inverso Amylin and the Stability of Amylin Fibrils. J. Chem. Theory Comput. 2020, 16 (8), 5358–5368. https://doi.org/10.1021/acs.jctc.0c00523.

(22) Gallardo, R.; Iadanza, M. G.; Xu, Y.; Heath, G. R.; Foster, R.; Radford, S. E.; Ranson, N. A. Fibril Structures of Diabetes-Related Amylin Variants Reveal a Basis for Surface-Templated Assembly. Nat. Struct. Mol. Biol. 2020, 27 (11), 1048–1056. https://doi.org/10.1038/s41594-020-0496-3.

(23) Kryshtafovych, A.; Moult, J.; Billings, W. M.; Della Corte, D.; Fidelis, K.; Kwon, S.; Olechnovič, K.; Seok, C.; Venclovas, Č.; Won, J.; CASP-COVID participants. Modeling SARS-CoV-2 Proteins in the CASP-commons Experiment. Proteins Struct. Funct. Bioinforma. 2021, 89 (12), 1987–1996. https://doi.org/10.1002/prot.26231.

(24) Mesias, V. St. D.; Zhu, H.; Tang, X.; Dai, X.; Liu, W.; Guo, Y.; Huang, J. Moderate Binding between Two SARS-CoV-2 Protein Segments and α-Synuclein Alters Its Toxic Oligomerization Propensity Differently. J. Phys. Chem. Lett. 2022, 13 (45), 10642–10648. https://doi.org/10.1021/acs.jpclett.2c02278.

(25) Abraham, M. J.; Murtola, T.; Schulz, R.; Páll, S.; Smith, J. C.; Hess, B.; Lindahl, E. GROMACS: High Performance Molecular Simulations through Multi-Level Parallelism from Laptops to Supercomputers. SoftwareX 2015, 1–2, 19–25. https://doi.org/10.1016/j.softx.2015.06.001.

(26) Huang, J.; Rauscher, S.; Nawrocki, G.; Ran, T.; Feig, M.; de Groot, B. L.; Grubmüller, H.; MacKerell, A. D. CHARMM36m: An Improved Force Field for Folded and Intrinsically Disordered Proteins. Nat. Methods 2017, 14 (1), 71–73. https://doi.org/10.1038/nmeth.4067.

(27) Jorgensen, W. L.; Chandrasekhar, J.; Madura, J. D.; Impey, R. W.; Klein, M. L. Comparison of Simple Potential Functions for Simulating Liquid Water. J. Chem. Phys. 1983, 79 (2), 926–935. https://doi.org/10.1063/1.445869.

(28) Wang, W.; Hansmann, U. H. E. Stability of Human Serum Amyloid A Fibrils. J. Phys. Chem. B 2020, 124 (47), 10708–10717. https://doi.org/10.1021/acs.jpcb.0c08280.

(29) Siwy, C. M.; Lockhart, C.; Klimov, D. K. Is the Conformational Ensemble of Alzheimer’s Aβ10-40 Peptide Force Field Dependent? PLOS Comput. Biol. 2017, 13 (1), e1005314. https://doi.org/10.1371/journal.pcbi.1005314.

(30) Samantray, S.; Yin, F.; Kav, B.; Strodel, B. Different Force Fields Give Rise to Different Amyloid Aggregation Pathways in Molecular Dynamics Simulations. J. Chem. Inf. Model. 2020, 60 (12), 6462–6475. https://doi.org/10.1021/acs.jcim.0c01063.

(31) Man, V. H.; He, X.; Derreumaux, P.; Ji, B.; Xie, X.-Q.; Nguyen, P. H.; Wang, J. Effects of All-Atom Molecular Mechanics Force Fields on Amyloid Peptide Assembly: The Case of Aβ _16–22_ Dimer. J. Chem. Theory Comput. 2019, 15 (2), 1440–1452. https://doi.org/10.1021/acs.jctc.8b01107.

(32) Bussi, G.; Donadio, D.; Parrinello, M. Canonical Sampling through Velocity Rescaling. J. Chem. Phys. 2007, 126 (1), 014101. https://doi.org/10.1063/1.2408420.

(33) Parrinello, M.; Rahman, A. Polymorphic Transitions in Single Crystals: A New Molecular Dynamics Method. J. Appl. Phys. 1981, 52 (12), 7182–7190. https://doi.org/10.1063/1.328693.

(34) Miyamoto, S.; Kollman, P. A. Settle: An Analytical Version of the SHAKE and RATTLE Algorithm for Rigid Water Models. J. Comput. Chem. 1992, 13 (8), 952–962. https://doi.org/10.1002/jcc.540130805.

(35) Hess, B.; Bekker, H.; Berendsen, H. J. C.; Fraaije, J. G. E. M. LINCS: A Linear Constraint Solver for Molecular Simulations. J. Comput. Chem. 1997, 18 (12), 1463–1472. https://doi.org/10.1002/(SICI)1096-987X(199709)18:12<1463::AID-JCC4>3.0.CO;2-H.

(36) Humphrey, W.; Dalke, A.; Schulten, K. VMD: Visual Molecular Dynamics. J. Mol. Graph. 1996, 14 (1), 33–38. https://doi.org/10.1016/0263-7855(96)00018-5.

(37) McGibbon, R. T.; Beauchamp, K. A.; Harrigan, M. P.; Klein, C.; Swails, J. M.; Hernández, C. X.; Schwantes, C. R.; Wang, L.-P.; Lane, T. J.; Pande, V. S. MDTraj: A Modern Open Library for the Analysis of Molecular Dynamics Trajectories. Biophys. J. 2015, 109 (8), 1528– 1532. https://doi.org/10.1016/j.bpj.2015.08.015.

(38) Pettersen, E. F.; Goddard, T. D.; Huang, C. C.; Couch, G. S.; Greenblatt, D. M.; Meng, E. C.; Ferrin, T. E. UCSF Chimera?A Visualization System for Exploratory Research and Analysis. J. Comput. Chem. 2004, 25 (13), 1605–1612. https://doi.org/10.1002/jcc.20084.

(39) Frishman, D.; Argos, P. Knowledge-Based Protein Secondary Structure Assignment. Proteins Struct. Funct. Genet. 1995, 23 (4), 566–579. https://doi.org/10.1002/prot.340230412.

(40) Kumari, R.; Kumar, R.; Open Source Drug Discovery Consortium; Lynn, A. G_mmpbsa —A GROMACS Tool for High-Throughput MM-PBSA Calculations. J. Chem. Inf. Model. 2014, 54 (7), 1951–1962. https://doi.org/10.1021/ci500020m.

(41) Bellaiche, M. M. J.; Best, R. B. Molecular Determinants of Aβ _42_ Adsorption to Amyloid Fibril Surfaces. J. Phys. Chem. Lett. 2018, 9 (22), 6437–6443. https://doi.org/10.1021/acs.jpclett.8b02375.

(42) Leguizamon Herrera, V. L.; Buell, A. K.; Willbold, D.; Barz, B. Interaction of Therapeutic D -Peptides with Aβ42 Monomers, Thermodynamics, and Binding Analysis. ACS Chem. Neurosci. 2022, 13 (11), 1638–1650. https://doi.org/10.1021/acschemneuro.2c00102.

