## Supplemental Figures and tables for "Human Amylin in the Presence of SARS-COV-2 Protein Fragments"

**Supporting information contains nine supporting figures (SF1-SF9) and one supporting table (ST1).**

**Table ST1**: Residue-wise binding frequencies of SK9 and FI10 residues to amylin monomers

| **SK9 (Autodock Vina) Binding Frequencies (%)** | | | | | | | | | |
|---|---|---|---|---|---|---|---|---|---|
| Residues | Ser-1 | Phe-2 | Tyr-3 | Val-4 | Tyr-5 | Ser-6 | Arg-7 | Ser-8 | Lys-9 |
| Last 800ns | 37 (6) | 63 (9) | 67 (4) | 70 (10) | 80 (10) | 55 (6) | 56 (2) | 50 (5) | 42 (3) |
| Last 200ns | 28 (3) | 58 (5) | 70 (20) | 70 (20) | 70 (20) | 50 (10) | 48 (8) | 30 (20) | 28 (9) |
| **SK9 (HADDOCK) Binding Frequencies (%)** | | | | | | | | | |
| Last 800ns | 41 (8) | 70 (10) | 70 (10) | 60 (20) | 80 (20) | 50 (10) | 60 (20) | 40 (10) | 30 (10) |
| Last 200ns | 40 (30) | 60 (20) | 60 (30) | 50 (30) | 60 (30) | 30 (10) | 50 (30) | 30 (10) | 30 (20) |

| **FI10 Binding Frequencies (%)** | | | | | | | | | | |
|---|---|---|---|---|---|---|---|---|---|---|
| Residues | Phe-1 | Lys-2 | Asn-3 | Ile-4 | Asp-5 | Gly-6 | Tyr-7 | Phe-8 | Lys-9 | Ile-10 |
| Last 800ns | 64 (5) | 54 (6) | 48 (7) | 76 (8) | 53 (6) | 50 (10 | 80 (4) | 86 (6) | 58 (5) | 77 (9) |
| Last 200ns | 70 (20) | 70 (20) | 54 (7) | 90 (10) | 60 (10) | 40 (30) | 91 (4) | 95 (3) | 60 (10) | 80 (20) |

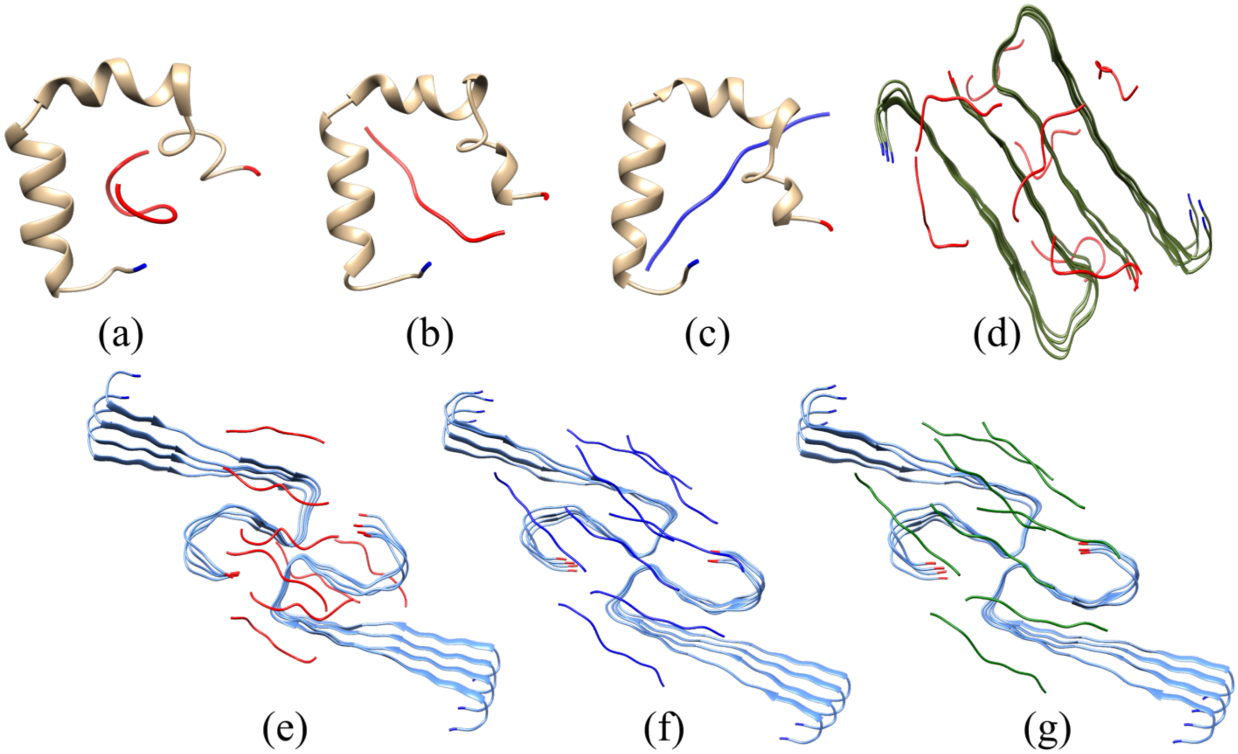


**Figure SF1**: Start configurations of amylin monomer (a-c), the 2F4L fibril (d) and the 6ZRV fibril (e-g) with SK9 (red) and FI10 (blue). N-termini are marked in dark blue and C-termini in red.

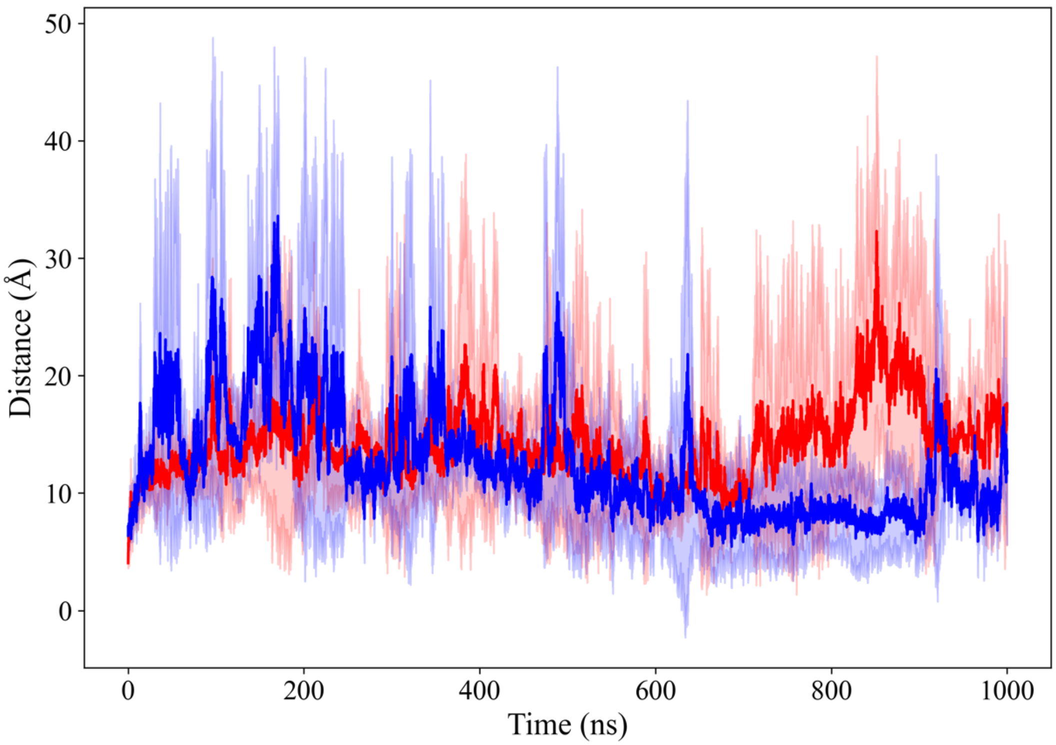


**Figure SF2**: Average center of mass distances as function of time between the amylin monomer and SK9 (red) or FI10 (blue)

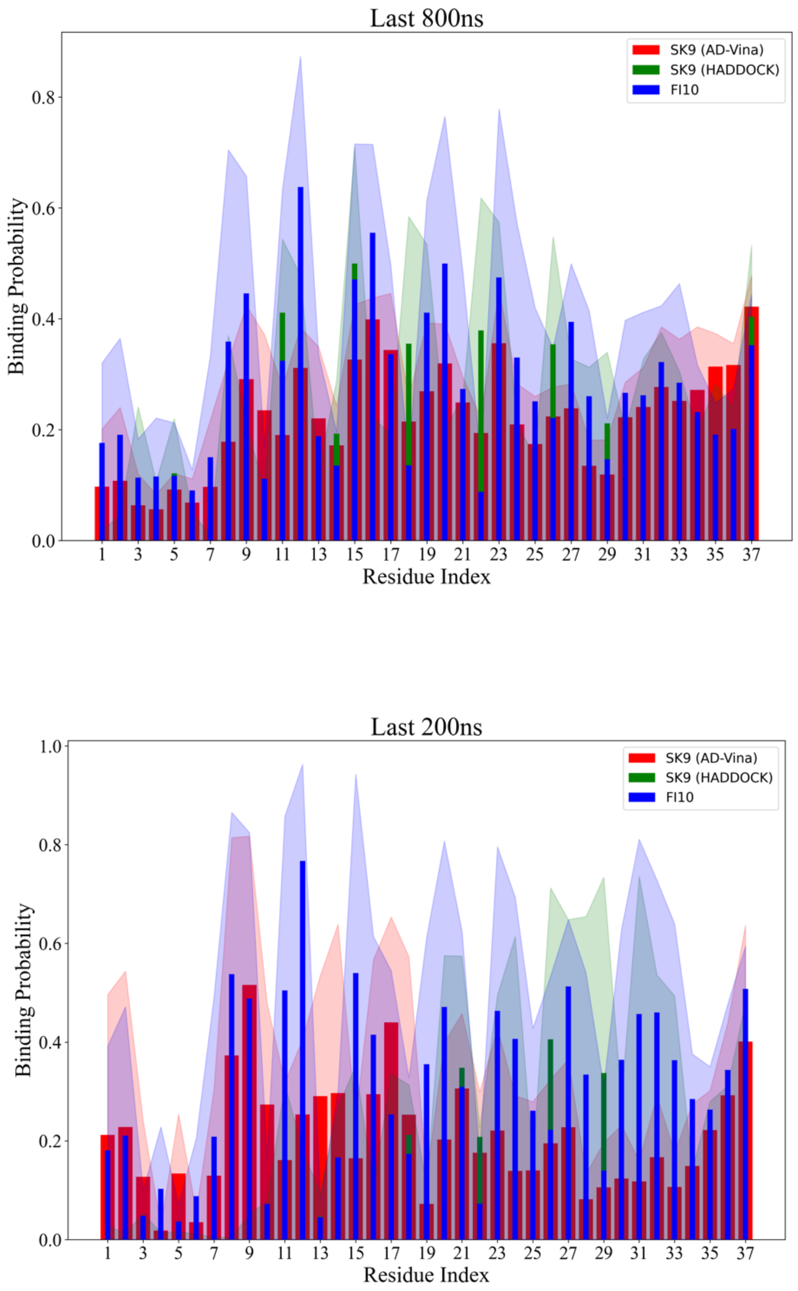


**Figure SF3**: Residue-wise binding frequencies for the amylin monomer interacting with SK9 or FI10 (blue). Binding sites for SK9 were found by two docking programs, Autodock Vina (red) and Haddock (green).

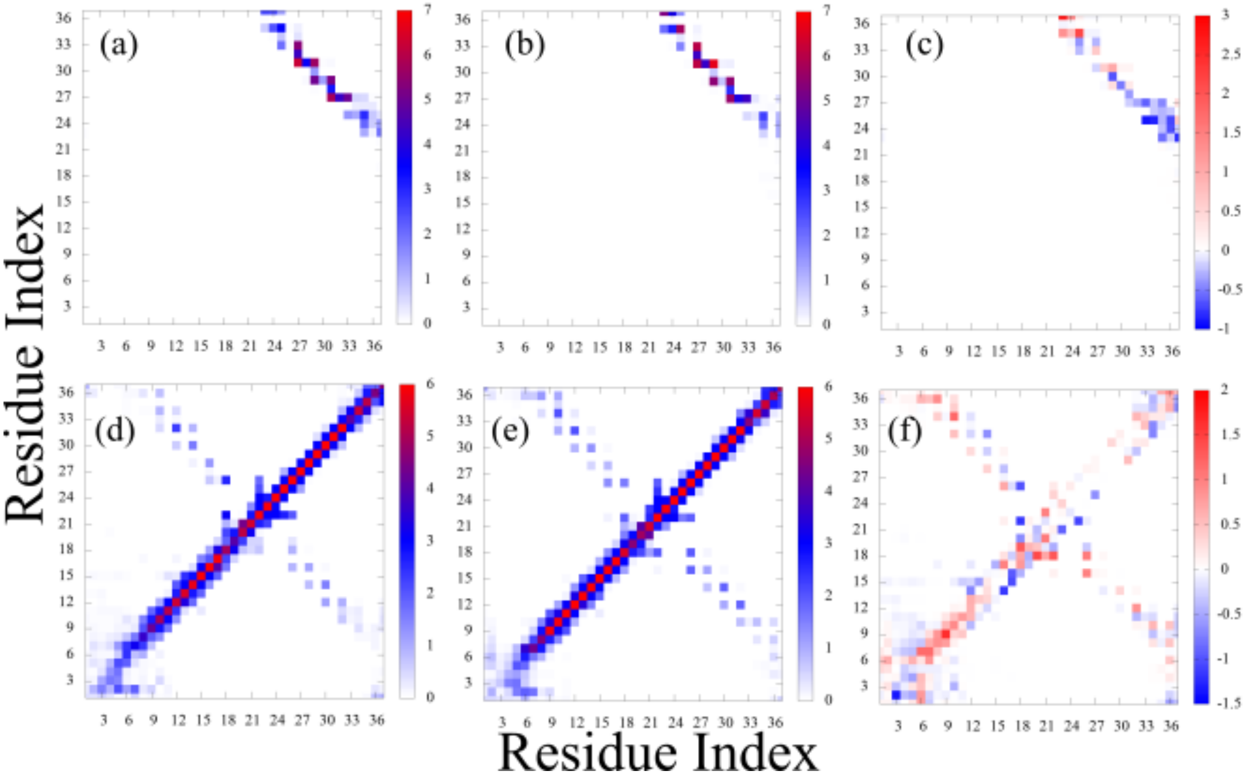


**Figure SF4**: Map of packing contacts, i.e., contacts between residues located on chains on different protofibrils for the (a) control and (b) in presence of SK9 of the fibril model 2F4L. The average number of such contacts are shown. The difference between the two systems is shown in (c). The corresponding maps for contacts between residues on chains on different layers, i.e., stacking contacts, are shown in (d)-(f).

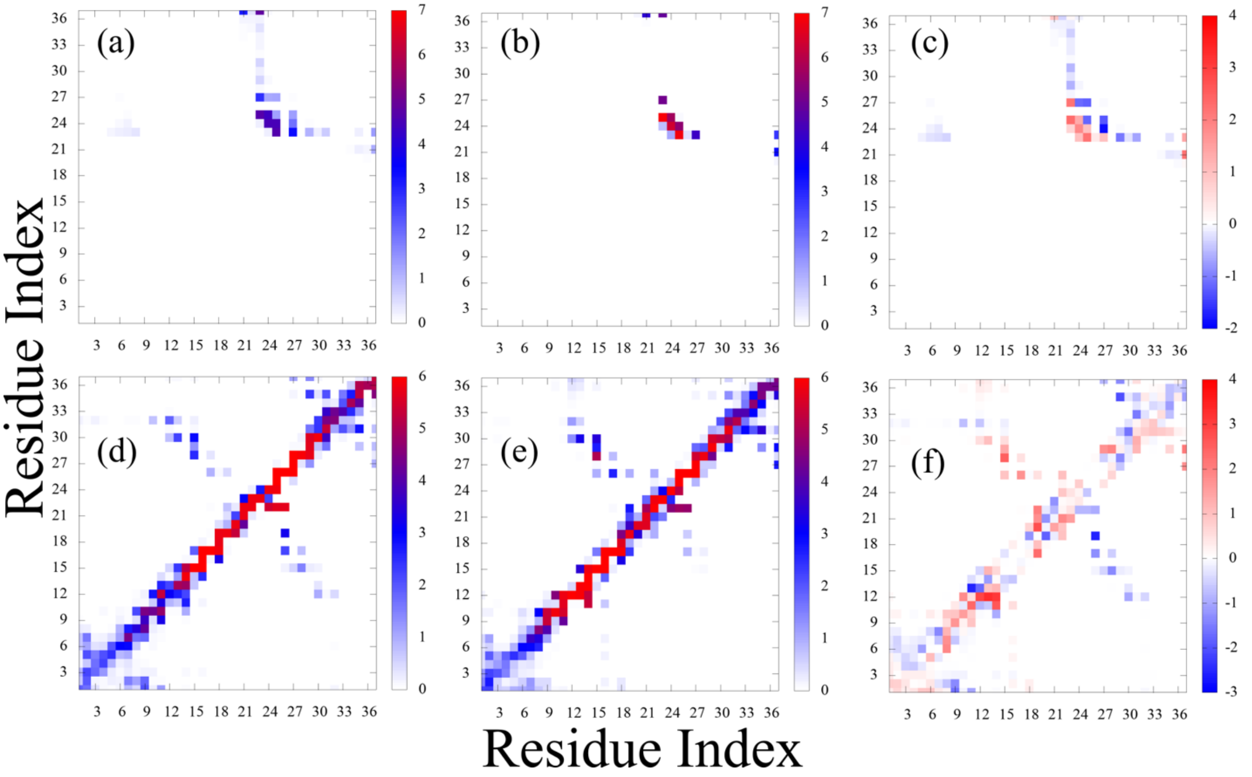


**Figure SF5**: Map of packing contacts, i.e., contacts between residues located on chains on different protofibrils in the fibril model with PDB-ID 6ZRF. Shown are maps for the (a) control and (b) in presence of SK9. Shown are the average number of such contacts. The difference between the two systems is shown in (c). The corresponding maps for contacts between residues on chains on different layers, i.e., stacking contacts, are shown in (d)-(f).

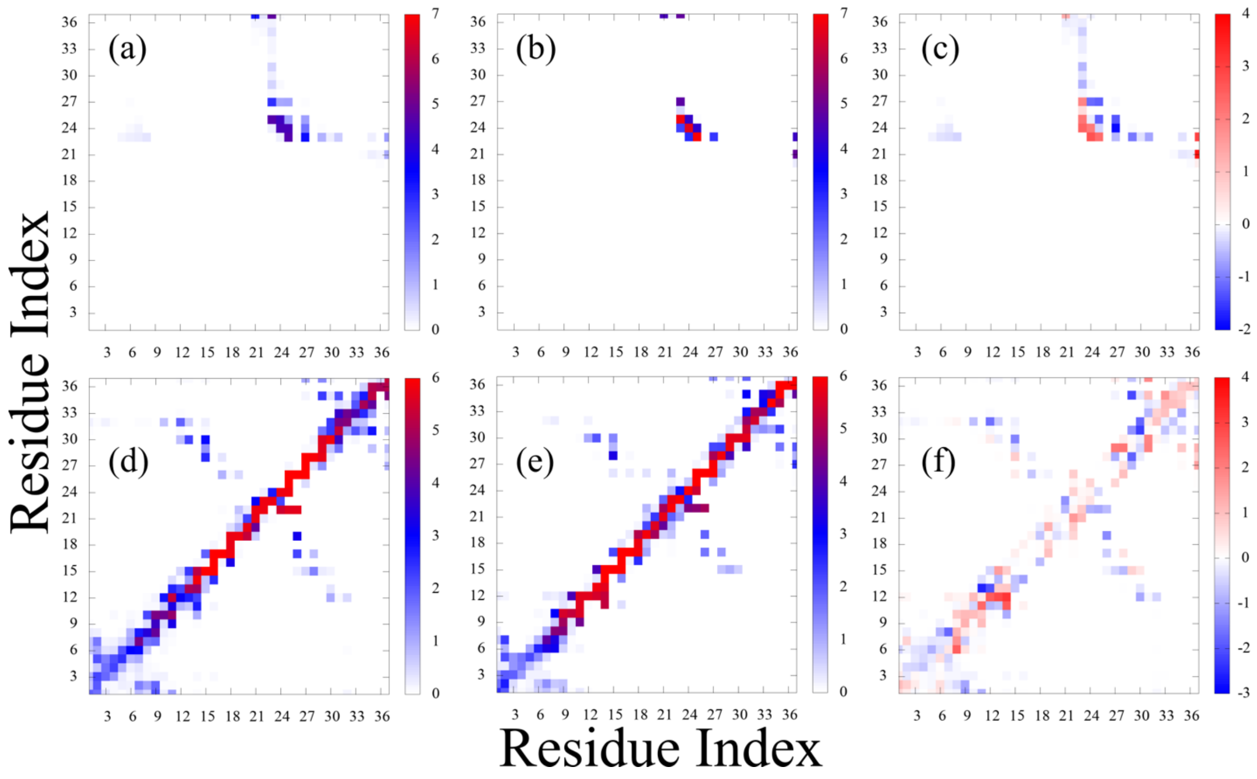


**Figure SF6**: Map of packing contacts, i.e., contacts between residues located on chains on different protofibrils in the fibril model with PDB-ID 6ZRF. Shown are maps for the (a) control and (b) in presence of FI10. Shown are the average number of such contacts. The difference between the two systems is shown in (c). The corresponding maps for contacts between residues on chains on different layers, i.e., stacking contacts, are shown in (d)-(f).

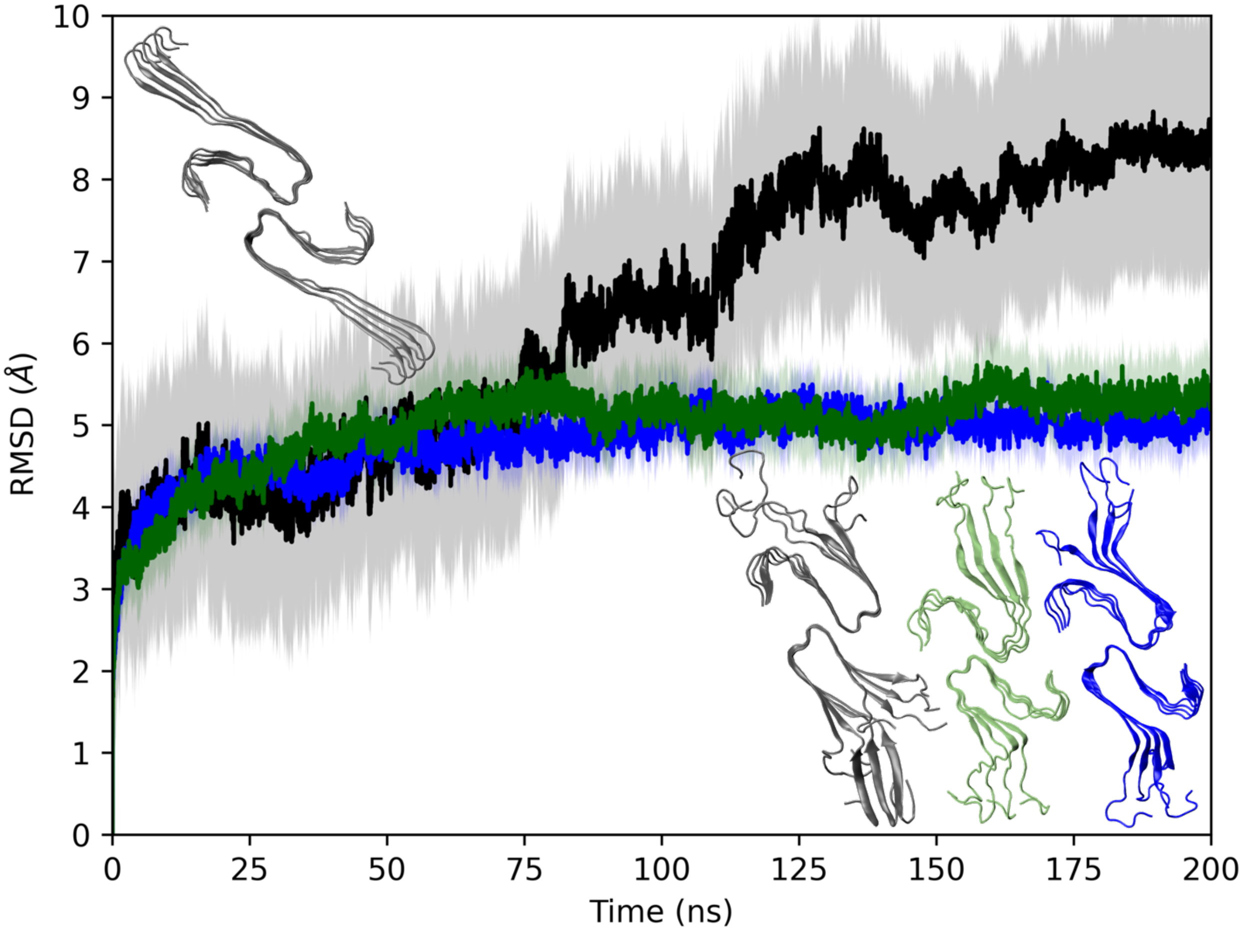


**Figure SF7**: RMSD to the start configuration as function of time for the fibril model 6ZRV. The RMSD is calculated over all backbone atoms in residues 8-37 of all chains in the respective fibril model. Black curves are from the control simulations while green curves are from data measured in simulations where FI10 is present and has a C-terminal end group COO-. Similarly, blue curves mark the case of FI10 with end group $CONH_2$ interacting with the amylin fibril. The standard deviation of the averages is shown shaded. The start configuration (top/left) and representative final configurations (right/bottom) of the amylin fibril models are shown as insets.

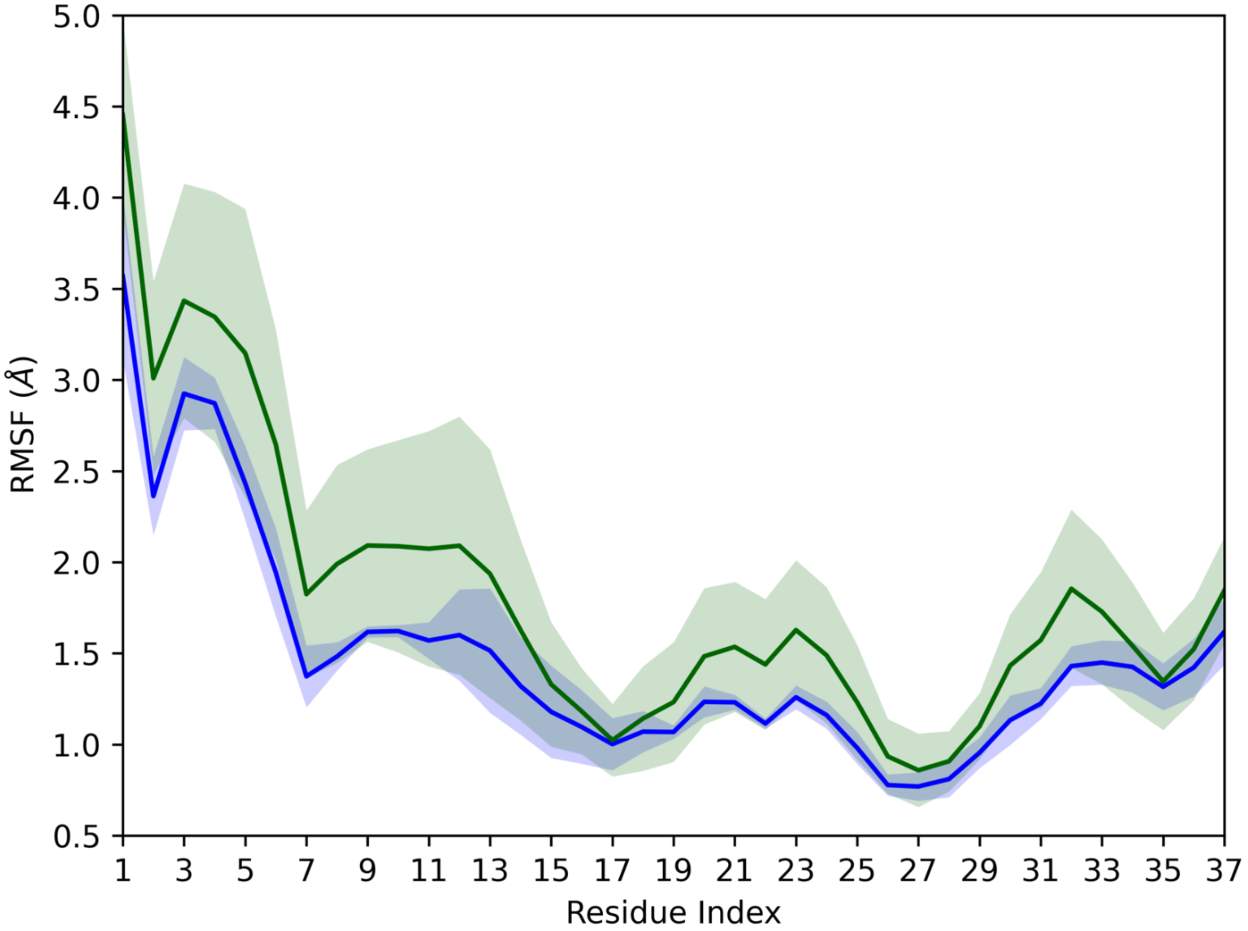


**Figure SF8:** Residue-wise RMSF for the fibril model 6ZRF in presence of FI10 with COO- (green) and $CONH_2$ (blue) end group. Shaded regions mark the standard deviation of the averages.

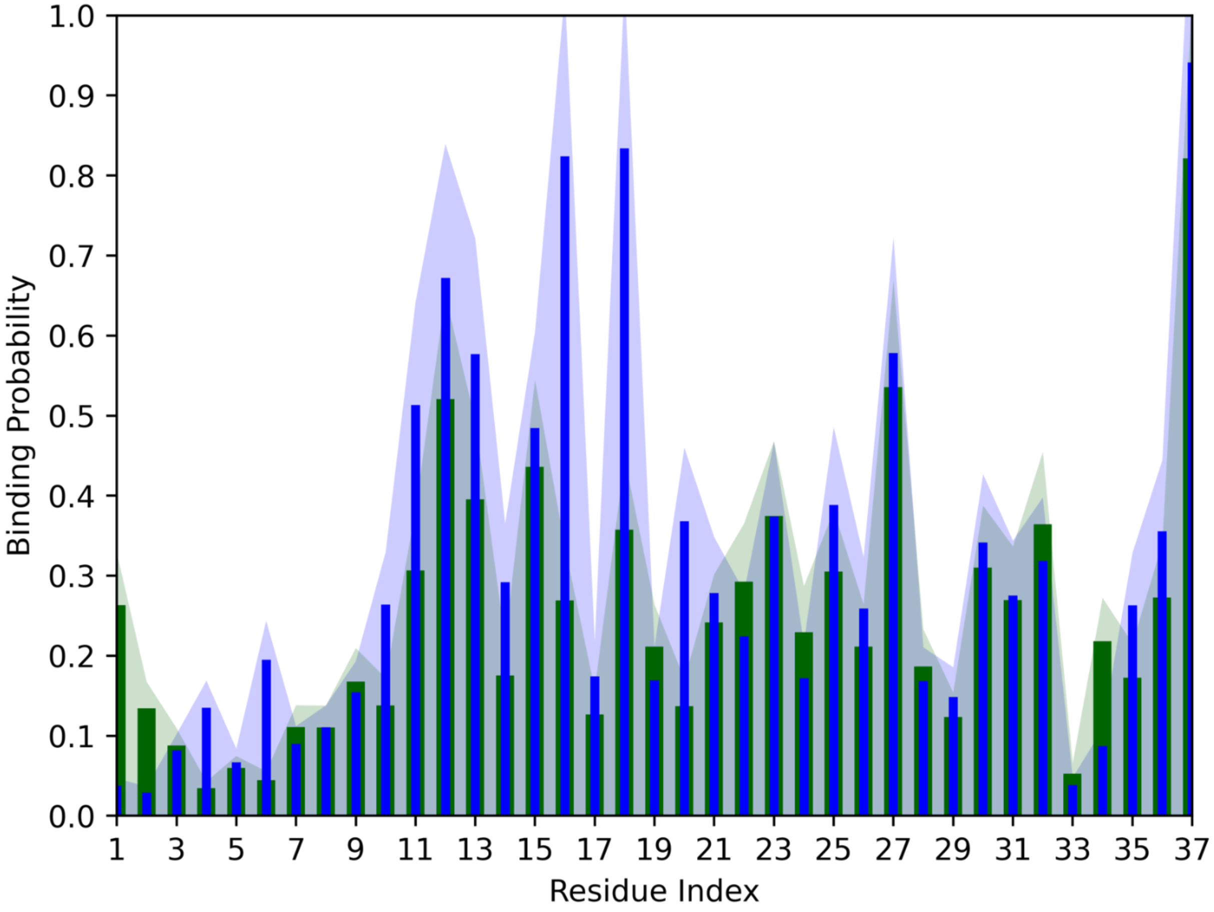


**Figure SF9**: Residue-wise frequency of contacts for the fibril model 6ZRF with FI10 having either COO- (green) or $CONH_2$ (blue) end group. Shaded regions mark the standard deviation of the averages.
