## Supplementary material for "Human Amylin in the Presence of SARS-COV-2 Protein Fragments": Folder with start and final configurations for all trajectories: README

Amylin Monomer and Fibril Start and Final Configurations:

All amylin monomer configurations can be found under CONFIGURATIONS-AMYLIN-MONOMERS/ where they are further seperated into Control, SK9, and FI10 simulations. SK9 contains two sets of start and final configurations. One for the simulations that used AutoDock-Vina for docking and the other for docking performed by HADDOCK.

Similarly the fibril configurations are located under CONFIGURATIONS-AMYLIN-FIBRILS/ which is further seperated into two fibril models utlized, 2F4L and 6ZRF. 

SARS-COV-2 Protein Fragments, SK9 and FI10, Configurations are located under SARS-COV-2-PROTEIN-FRAGMENTS
